# Remodeling of KRAS interactome induced by clinically relevant RAS inhibitors reveals convergent responses and a KRAS-mediated regulation of directional cell migration

**DOI:** 10.1101/2025.11.14.688534

**Authors:** Yang Yang, Charles Y. Cho, Ansgar Brock, Doug Quackenbush, Kaisheng Chen, Frederick Lo, Cyril H. Benes, Jacob R. Haling

## Abstract

KRAS is commonly mutated in lung, colorectal, and pancreatic cancers. Small molecule inhibitors targeting KRAS with distinct mechanisms-of-action and variable specificities have entered the clinic, but a comprehensive view of their effect on the RAS signaling network has not been reported. Here, we describe the impact of RAS inhibition on the KRAS protein interactome using Proximity-dependent Biotinylation Identification. Two inhibitors were used: panRAS-ON, which forms a ternary complex between cyclophilin and RAS in the GTP bound state; and panKRAS-off, which binds specifically to the KRAS switch II pocket. RAS inhibitors were found to significantly alter 16.5% of proteins in proximity to KRAS. Despite their distinct mechanisms-of-action, the KRAS inhibitors induced highly correlated changes in the KRAS interactomes. Among proteins in close proximity to KRAS, Afadin was found to be highly regulated by RAS inhibition. AFDN has been characterized as a key regulator of cell motility, invasion, and metastasis. Analysis of AFDN phosphorylation revealed that AKT only partially modulates p-Ser1718, while inhibition of KRAS is sufficient to abolish EGF-mediated AFDN phosphorylation. Knockdown of AFDN in a KRAS-driven non-small cell lung cancer model abolished chemotaxis in a transwell migration assay and disrupted directional movement in an EGF-driven wound healing model. These results suggest that KRAS is a central node in regulating growth factor induced cell migration, and that KRAS inhibition plays a broader role than MAPK-mediated cell proliferation and survival.

## Introduction

KRAS is a key signaling protein that transmits extracellular growth signals to downstream pathways by recruiting a diverse array of effector proteins. The widely accepted model of KRAS signaling involves the active, GTP-bound KRAS directly interacting with effector proteins such as the RAF kinases and PI3K, thereby triggering downstream signaling pathways, including the mitogen-activated protein kinase (MAPK) and phosphatidylinositol 3-kinase (PI3K) pathways (1). As a small GTPase, KRAS possesses intrinsic GTP hydrolysis activity, which, along with the regulatory actions of GTPase activating proteins (GAPs), facilitates the hydrolysis of GTP, converting activated KRAS into its inactive, GDP-bound form (1,2). This negative regulation ensures that the KRAS-mediated signaling cascade in normal cells is precisely controlled, maintaining the critical role of KRAS in cell proliferation and survival within appropriate limits and timeframes.

KRAS is among the most frequently mutated oncogenes, with a mutation rate of 31.2% in all human cancers, especially in the three most lethal cancer types (78.7% in pancreatic cancer, 49.7% in colorectal cancer, 20.1% in lung cancer) (3). Mutant KRAS remains predominantly in the active GTP-bound state due to impaired intrinsic GTPase function and reduced sensitivity to negative regulation by GAPs, leading to constitutive activation of pro-neoplastic signaling and driving malignant transformation (3,4). Although once deemed “undruggable”, KRAS now has emerged as a viable drug target with two FDA-approved therapies directly targeting KRAS^G12C^ and several others in various stages of clinical development (5,6). Despite demonstrated clinical benefit, many patients have de novo or acquired resistance to single-agent treatment (7). A more complete understanding of KRAS signaling and its interactome could help identify new therapeutic targets, suggest combination strategies, and understand possible resistance mechanisms in KRAS-driven cancers.

Decades of research have expanded the KRAS-regulated signaling networks beyond the canonical RAF/MAPK or PI3K pathways to include a variety of newly discovered interactors, such as LZTR1 (8), PIP5K1A (9), mTORC2 (10), and others (11). Discovery of these novel interactors has been substantially driven by advanced interactome screening techniques. In particular, the application of Proximity Labeling-coupled Mass Spectrometry (PL-MS) to investigate the KRAS interactome starting from 2018 has greatly expanded the number of KRAS-associated protein interactions (Fig. S1A). To date, interactions identified by PL-MS account for 72.1% of the total 1951 published KRAS biophysical interactions (Fig. 1A). While traditional methods like yeast-hybrid or affinity purification rely on extracting stable binary protein interactions in cell extracts, they often fail to capture the transient and dynamic protein interactions integral to KRAS signaling. PL-MS, however, represents an alternative approach that enables high-throughput and unbiased chemoproteomic profiling of local protein interactions in living cells (12,13).

**Figure. 1.**
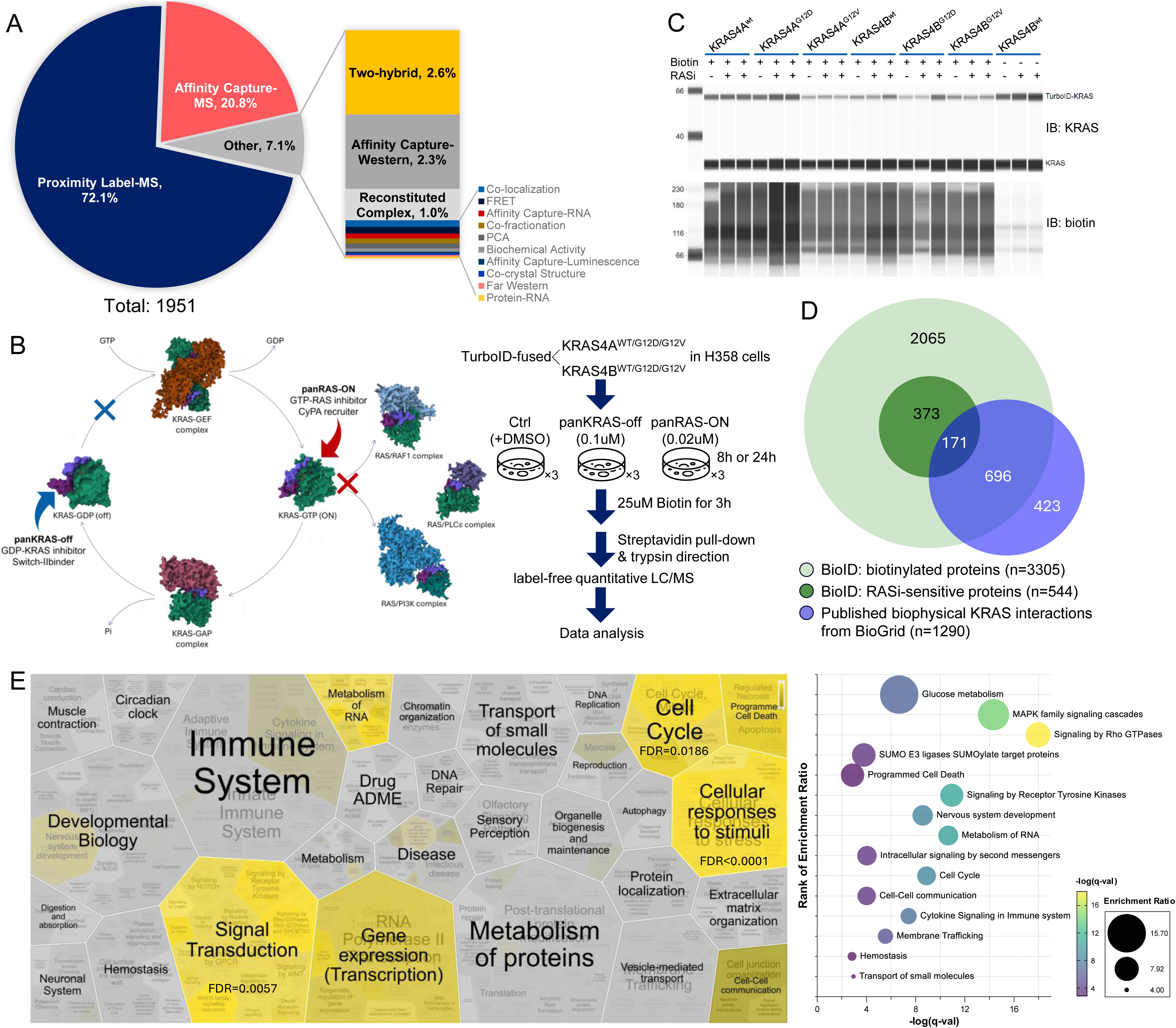
BioID efficiently identifies KRAS interactomes influenced by pharmacological perturbation (A) Published KRAS biophysical interactions categorized by experimental approach from BioGrid. (B) Schematic of mechanisms of action for the two selected RAS inhibitors and experiment workflow of the BioID study. H358 cells expressing different KRAS variants were treated with panRAS-ON or panKRAS-off for 8 hours or 24 hours with 25 µM Biotin added to the cells 3 hours before harvesting. LC/MS, liquid chromatography-tandem mass spectrometry. KRAS is shown in green, with switches I and II colored differently. PDB IDs: KRAS-GDP: 4BOE; KRAS-GTP: 5UK9; KRAS-GEF: 7KFZ; KRAS-GAP: 6OB2; KRAS-RAF: 6XHB; KRAS-RGL1: 7SCW; KRAS-PI3K: 1HE8. GEF: guanine nucleotide exchange factor. (C) Immunoblot of TurboID-fused KRAS variants (top) and promiscuously biotinylated proteins (bottom) in lentiviral-transduced H358 cells with (+) or without (-) biotin addition. Endogenous KRAS are shown to compare expression levels and as loading controls. (D) Overlap of biotinylated proteins from our BioID study (light green, n=3305) with published KRAS interactors from BioGrid (blue, n=1290). The subset of 544 KRAS-proximal proteins significantly affected by RAS inhibitors among total biotinylated proteins were highlighted in dark green (log2FC>1 or <-1, FDR<0.05) (E) Enrichment analysis of top Reactome pathways from the subset of 544 KRAS interactor candidates using Reactome analysis tool (https://reactome.org/PathwayBrowser/#TOOL=AT). Major category of pathways overrepresented among the subset of 544 KRAS interactors were highlighted in yellow on the left. The detailed top 15 significantly enriched Reactome pathways were listed on the right. Fill color represents the significance of enrichment in the KRAS interactome, and size represents the enrichment ratio of each pathway in the KRAS interactome

Our proximity-dependent biotin identification (BioID) methodology incorporates the biotin ligase TurboID, which promiscuously conjugates biotin to proteins within a 10nM radius. These biotinylated proteins can then be captured and analyzed via liquid chromatography-mass spectrometry (LC-MS) (13). In this study, we utilized two clinically relevant RAS inhibitors (RASi) with distinct mechanisms of action (MOA) to inhibit KRAS in cancer cells and evaluate the changes in KRAS interactome using BioID. Profiling of KRAS proximity revealed that RASi targeting different nucleotide states of KRAS through distinct MOA displaced nearly identical KRAS interactors, suggesting accumulation of the inactive GDP-bound KRAS can be functionally equivalent to CYPA-mediated direct inhibition of active GTP-bound KRAS. Among the KRAS interactors identified with high confidence, Afadin (AFDN) emerged as a top hit, consistently disrupted by RASi from KRAS proximity across all variants. Subsequent biological analyses revealed that the EGF-induced phosphorylation of AFDN is primarily mediated by KRAS activity. Furthermore, AFDN-dependent directional cell migration was markedly suppressed by RASi. These results highlight how proteomic network analysis in combination with pharmacological inhibition of key nodes can help further understanding of oncogenic pathways.

## Methods

### Construction of KRAS-TurboID stable cell lines

For BioID experiments, the constructs encoding 3×FLAG-TurboID fused to the N-terminus of KRAS4A (NP_203524.1), KRAS4B (NP_004976.2) and mutants were created by gene synthesis and cloned into the pCDH-EF1-CMV-Puro plasmid, leading to generation of the following vectors: pCDH PuroR-TurboID KRAS4A^wt^, pCDH PuroR-TurboID KRAS4A^G12D^, pCDH PuroR-TurboID KRAS4A^G12V^, pCDH PuroR-TurboID KRAS4B^wt^, pCDH PuroR-TurboID KRAS4B^G12D^, and pCDH PuroR-TurboID KRAS4B^G12V^. The sequence<u>s</u> of synthesized cDNA are listed in supplementary <u>file</u> 1. All constructs were verified by sequencing. Lentiviruses of each vector were produced by co-transfection of vectors with Delta8.9 and pVSV-G in HEK293T cells. For stable cell lines, NCI-H358 cells were seeded at 1×10^6^ cells per well in 6-well plates in Dulbecco’s Modified Eagle’s Medium containing 10% fetal bovine serum. Lentivirus were added into each well followed by polybrene to a final concentration of 8 µg/ml. At 48 hours, transduced cells were selected with 1 µg/ml puromycin until all cells in the control well were depleted.

### Biotin Identification

BioID was performed in human non-small cell lung cancer H358 cells stably expressing the six different KRAS variants (wild-type KRAS4A, KRAS4A^G12D^, KRAS4A^G12V^, wild-type KRAS4B, KRAS4B^G12D^, KRAS4B^G12V^) with FLAG-tagged TurboID fused to the N-terminus of KRAS. The six derived cultures were propagated and grown to 80% confluency in 15 cm tissue culture dishes followed by treatment with 0.02 µM panRAS-ON (Pan-RAS-IN-7, CAS No. 2642135-72-4),

0.1 µM panKRAS-off (Pan KRas-IN-1, CAS No. 2791263-84-6) or DMSO in triplicate for 8 hours and 24 hours, respectively. Biotin was added to each plate to a working concentration of 25 µM in the last 3 hours of incubation. Harvested cells were lysed in pre-chilled lysis buffer (20 mM HEPES pH 8.0, 150 mM NaCl, 1% NP40, Halt protease & phosphatase inhibitor (Thermo Fisher, Cat # 78444), and Pierce Universal Nuclease) by end-to-end rotation at 4℃ for 1 hour. Lysates were centrifuged at 14,000 rpm for 15 minutes and the supernatants were incubated with Pierce Streptavidin Magnetic Beads (Thermo Fisher, Cat # 88816) by end-to-end rotation at room temperature for 1 hour. After binding, the collected beads were washed twice with 1% SDS (in 100 mM Tris, pH 8.0), 5 times with 8M Urea (in 100 mM Tris, pH 8.0), and 5 times with 20% Acetonitrile (ACN). Next, the beads were resuspended in buffer (100 mM Triethylammonium bicarbonate (TEAB) and 10% ACN) and digested with trypsin at 37℃ overnight. The following day, the beads were removed and the supernatants containing digested peptides were air dried and resuspended in 0.1% formic acid for mass spectrometry analysis.

### Proteomics Data Acquisition

Solid-phase Extraction purified on-bead digests were loaded onto Evotips (EV2018, Evosep) using a modified protocol. Evotips were loaded with 20 µL of DMAc followed by spinning at 500 g for 1 minute. This was followed by loading tips with 200 µl of 0.1% formic acid buffer and spinning at 500 g for 1 minute. Samples were added to the remaining liquid in the tips followed by spinning at 500 g for 2 minutes. Extra liquid in the tips samples were spun repeatedly for 1 minute intervals at 500rcf until no liquid remained in the tips. Tips were then loaded with 200 µl of 0.1% formic acid buffer and spun for 1 minute at 500 g before being either stored at 4℃ or placed onto the instrument for acquisition.

Peptides were separated using an Evosep One liquid chromatography system employing the 15 samples per day method. A 150 mm long, 100 µm ID reversed-phase column with 3 µm particles (#1895806, Bruker) was used. Separation was performed at room temperature. The LC was coupled to an Orbitrap Fusion Lumos Tribrid mass spectrometer (Thermo Fisher Scientific) via a Nanospray Flex ion source (ES071, Thermo Fisher Scientific) using a Nanospray Flex adapter (EV1085, Evosep) and stainless-steel emitter (EV1086, Evosep).

MS data were acquired in data-dependent acquisition (DDA) mode. Full MS scans were acquired in the Orbitrap at a resolution of 60,000 over a mass range of 375–1500 m/z. The automatic gain control (AGC) target was set to Standard with a maximum injection time of 50 milliseconds. A HCD normalized collision energy of 30% was used and MS/MS spectra were acquired in the Orbitrap at a resolution of 15,000 with an AGC target of 5 × 10⁴ and a maximum injection time of 100 ms. Dynamic exclusion time was set to 30 s.

Raw data files were processed using MaxQuant (version 2.5.1) with default parameters. Spectra were searched against the UniProt human reference proteome database (downloaded 06202022, one protein sequence per gene). Trypsin/P was specified as the digestion enzyme, allowing up to two missed cleavages. Carbamidomethylation of cysteine was set as a fixed modification, while oxidation of methionine, Glu to pyro-Glu, and N-terminal protein acetylation were set as variable modifications. Precursor ion quantification was performed using default settings.

### Cell signaling immunoblot analysis

Parental H358 cells were lysed in cold Pierce IP Lysis Buffer (Thermo Fisher, Cat # 87787) with Halt™ Protease and Phosphatase Inhibitor Cocktail (Thermo Fisher, Cat # 78440). Crude lysates were centrifuged at 14,000 rpm for 15 minutes and the total protein concentration of supernatant was determined by Pierce™ BCA Protein Assay Kits (Thermo Fisher, Cat # 23227). One microgram of total protein per sample was loaded on the Jess Automated Western Blot System (Bio-Techne) using the following primary antibodies: K-Ras Recombinant Rabbit Monoclonal Antibody (11H35L14) (Invitrogen, Cat # 703345); Anti-l/s-Afadin antibody produced in rabbit (Sigma, Cat # A0224); Phospho-Afadin (Ser1718) Antibody #5485 (Cell Signaling); Phospho-p44/42 MAPK (Erk1/2) (Thr202/Tyr204) (D13.14.4E) XP® Rabbit mAb #4370 (Cell Signaling); p44/42 MAPK (Erk1/2) Antibody #9102 (Cell Signaling); Phospho-Akt (Ser473) (D9E) XP® Rabbit mAb #4060 (Cell Signaling); Akt (pan) (C67E7) Rabbit mAb #4691 (Cell Signaling); GSK-3β (27C10) Rabbit mAb #9315 (Cell Signaling); Phospho-GSK-3-beta (Ser9) (D3A4) Rabbit mAb #9322 (Cell Signaling)

### Co-immunoprecipitation assays

H358 cells stably transduced with FLAG-tagged TurboID-fused KRAS4A^G12V^ or KRAS4B^G12V^ were grown to 80% confluency and treated with 0.02 µM panRAS-ON, 0.2 µM panKRAS-off, 0.1 µM Trametinib or DMSO for 1 hour. Next, cells were lysed in ice-cold buffer containing 20 mM Tris (pH 7.5), 150 mM NaCl, 1 mM DTT, 5 mM MgCl2, 0.5% NP-40, 0.5% Triton X-100, and HALT protease/phosphatase inhibitor for 5 minutes. Cell lysates were centrifuged at 14,000 rpm for 15 minutes and the total protein concentration from the supernatant was measured by BCA assay. 1.6 mg of total cell lysate were incubated with 40 µl anti-FLAG M2 beads (Sigma, Cat # M8823) for 40 minutes by end-to-end rotation at 4℃. The beads were then washed 3 times with 1x TBST to remove unbound proteins and eluted with 50 µl Elution buffer (200 µg/ml 3xFLAG peptide in lysis buffer). Lysate and eluate of each sample were collected for immunoblot on the Jess Automated Western Blot System.

### Generation of AFDN knockdown cells and rescues

*AFDN* knockdown H358 cell lines were constructed using lentiviral-shRNA purchased from Horizon Discovery: SMARTvector Lentiviral Human AFDN hCMV-None shRNA (Cat# V3SH7590-228832092, V3SH7590-228385407, V3SH7590-226768045); SMARTvector Non-targeting hCMV Control Particles (Cat#: VSC7078). Briefly, 1×10^6^ cells per well were seeded into 6-well plates in complete DMEM medium with 10% FBS. The next day, the cells were incubated with 1 ml fresh growth medium with 8 µg/ml polybrene for 1 hour followed by addition of lentivirus particles to MOI=2. 48 hours post transfection, medium was removed and replaced with selection media (1 µg/ml puromycin in DMEM with 10% FBS) until non-infected control cells were depleted. Knockdown efficiency was confirmed by immunoblotting.

To re-express the AFDN RA1/2 domain deletion constructs (RAΔAFDN), a synthetic DNA sequence encoding N-terminally His-tagged RAΔAFDN (aa350-1824 of P55196-4) cloned into pCMV6-AC vector were purchased from Eurofins Scientific (Supplementary file 1). The plasmid sequence was codon optimized to ensure resistance to silencing from shAFDN in the knockdown cell lines. shAFDN-H358 cells were transiently transfected with the RAΔAFDN construct using Lipofectamine™ 3000 Transfection Reagent (Thermo Fisher, Cat # L3000001) according to manufacturer’s instruction. Protein expression and phosphorylation of the rescue cell lines were tested 48 hours post transfection.

### EGF-stimulated AFDN phosphorylation

Parental H358 cells or the RAΔAFDN-rescued cells were seeded in 12-well plates in DMEM with 10% FBS and grown to 70% confluence. Next, the cells were serum-starved overnight followed by EGF stimulation carried out by replacing media with fresh serum-free DMEM containing 100 ng/ml recombinant human EGF protein (BioLegend, Cat# 585506). Cells were harvested at the indicated time intervals and lysed in RIPA Lysis and Extraction Buffer (Thermo Fisher, Cat # 89901) supplemented with Halt protease & phosphatase inhibitor added before use. Lysate protein concentration was determined by BCA assay, and 1 µg protein of each lysate was analyzed on the Jess Automated Western Blot System.

To investigate the impact of different kinase inhibitors on AFDN phosphorylation, H358 cells were seeded in 12-well plate and grown until 70% confluency. Next, the cells were serum-starved overnight and treated with different inhibitors for 1 hour at the following final concentrations: 0.02 µM panRAS-ON, 0.1 µM panKRAS-off, 0.1 µM adagrasib (KRAS^G12C^i), 0.1 µM trametinib, 1 µM MK2206 (pan-AKTi, AdipoGen Life Sciences, CAT# SYN-1162-M005) or DMSO as control. Final concentration of 10% FBS or 100 ng/ml EGF were immediately added to the plate for 5 minutes and the cells were harvested for immunoblotting.

### Cell motility

H358 cells were seeded into Revvity PhenoPlates (6057302) at the density of 2000 cells per well. After cell attachment, the wells were treated with 3-fold serial dilution of panRAS-ON (starting at 20 nM) and panKRAS-off (starting at 300 nM), respectively, in the presence of NucSpot Live650 (Biotium, Cat # 40082) for nuclear staining. The plates were incubated at 37℃ for 24 hours. High-content live-cell imaging was performed using a Phenix Plus system (Revvity) equipped with a spinning disk confocal module (50 µm pinhole) and an Andor Zyla 4.2 sCMOS camera (2160 × 2160 ROI, 2 × 2 binning). Samples were maintained at 37 °C, 5% CO₂, and 100% humidity.

Autofocus was achieved via a two-peak laser routine. Images were acquired with a 20× water-immersion Plan Fluor objective (NA 1.0), 40 ms exposure, using a 640 nm laser (140 mW, 25% power) for excitation and a 640–760 nm bandpass filter for emission. For each well, 4 fields of view were collected as single z-planes with 2% overlap. Time-lapse acquisition comprised 172 time points at 15-minute intervals.

### Transwell assay

The transwell assay was performed using Corning® 6.5 mm Transwell® with 8.0 µm Pore Polyester Membrane Insert (Corning, Cat # 3464) according to the manufacturer’s instructions. Briefly, cells were resuspended to 4×10^5^ cells/ml in serum-free RPMI media and 200 µl cell suspension was added to each transwell insert. 20% FBS in RPMI medium was added to the bottom well as chemoattractant, with 0.5% FBS as negative control, and cells were incubated at 37℃ for 60 hours. Inserts were washed with PBS followed by fixation using 4% paraformaldehyde and staining with 0.1% crystal violet.

### In vitro wound-healing assay and imaging acquisition

In vitro wound healing assay was performed with 2-well Culture-Insert in 24-well plates (Ibidi, Cat # 80242) according to manufacturer’s instructions. Stable shAFDN-knockdown H358 cells and the non-targeting control shRNA-transduced cells were grown to 60% confluency, trypsinized, and resuspended to 8×10^5^ cells/ml. 70 µl of the cell suspension were added into each chamber of the 2-well culture inserts. After the cells are completely confluent, cells were serum starved overnight and treated with RASi as required. Next day, the culture inserts were removed, and the media were replaced with fresh Phenol-free RPMI medium with NucSpot Live 650 (Biotium, Cat # 40082) for live cell staining. EGF was added into the wells to a final concentration of 100ng/ml. Images were captured every 16 minutes for a continuous 48 hours duration using a 10x objective on a Phenix High-Content Imaging System. High-content live-cell imaging was performed using a Phenix Plus system (Revvity) equipped with a spinning disk confocal module (50 µm pinhole) and an Andor Zyla 4.2 sCMOS camera (2160 × 2160 ROI, 1 × 1 binning). Autofocus was achieved via a two-peak laser routine. Images were acquired with a 10× Plan Fluor objective (NA 0.30), 200 ms exposure, using a 640 nm laser (140 mW, 50% power) for excitation and a 640–760 nm bandpass filter for emission. For each well, 36 fields of view were collected as single z-planes with 2% overlap. Time-lapse acquisition comprised 180 time points at 16-minute intervals Raw images generated from Phenix were background corrected and then segmented using Cellpose (14) to identify individual cells in each image. For individual cell tracking, both raw and segmented images are cropped to remove edge areas on both ends of the gap where cells move into. Then both raw and segmented image time-series were feed into ilastik (15), using its object tracking algorithm to track individual cell movement through the time course. The exported tracking results containing coordinates of each trackable cell across the entire time course are used to calculate a set of standard aggregated cell tracking features such as total distance, net distance, global and relative turning angle using definition and python scripts published by Svensson et al (16).

## Results

### BioID identifies a comprehensive KRAS interactome for both wild-type and oncogenic mutants

Proximity-dependent biotin identification (BioID) offers distinct advantages over conventional protein-protein interaction (PPI) screening methods by capturing weak or transient interactions in living cells. We used BioID to delineate the functional KRAS interactome through direct pharmacological inhibition of KRAS using two recently developed small molecule RAS inhibitors: panRAS-ON and panKRAS-off. BioID was performed in KRAS^G12C^ mutant H358 cells ectopically expressing TurboID fusions of wild-type KRAS or oncogenic mutants KRAS^G12D^ and KRAS^G12V^ for both splice variants KRAS4A and KRAS4B (Fig 1B). TurboID-fused KRAS proteins were expressed at moderate levels, slightly below endogenous KRAS (Fig 1C). Stable cell lines were treated with DMSO or either RASi for 8 or 24 hours, followed by biotin treatment, streptavidin pulldown, tryptic digestion, and mass spectrometry analysis. RASi did not alter native or TurboID-KRAS expression or promiscuous biotinylation in H358 cells (Fig 1C). We detected 3305 biotinylated proteins with an average of 2,376 proteins (2218 to 2563) per sample set with a substantially longer biotinylation time, greatly exceeding the size of interactome detected in previous studies using BirA-fusions (4). 867 biotinylated proteins overlapped with a published cohort of 1290 proteins known to have biophysical interactions with KRAS (67%, Fig 1D), indicating high coverage of the KRAS interactome.

Previous BioID studies focused on static interactions between KRAS4B and its effectors. We built a comprehensive molecular network for KRAS4A and KRAS4B whose interactions susceptible to two mechanistically distinct RASi in NSCLC cells (Fig 1B): panRAS-ON (Pan-RAS-IN-7) recruits the intracellular chaperone cyclophilin A (CYPA) to form a ternary complex with GTP-bound KRAS, HRAS, and NRAS, thereby inhibiting RAS function through occlusion of pathway activating PPIs; panKRAS-off (Pan KRas-IN-1) binds non-covalently to inactive GDP-bound KRAS, altering the conformation of the Switch I and II loops and preventing pathway activation. We compared the abundance of individual proteins in RASi-treated cells to the corresponding control group for all KRAS variants examined in the study (Fig. S1B). By removing proteins with less than a significant 2-fold change (FDR<0.05) between DMSO- and RASi-treated cells, we narrowed down the pool of candidate KRAS interactors to 544 proteins consistently displaced by RASi in the proximity of at least one KRAS variant (Fig. 1D). This allowed us to refine the functional KRAS interactome by focusing on proteins directly responding to pharmacological perturbation of KRAS while mitigating the challenge of promiscuous biotinylation associated with ectopic TurboID-KRAS.

Pathway enrichment analysis of the 544 RASi-responsive proteins revealed various biological processes affected by RASi, including cellular responses to stimuli, signal transduction, and cell cycle (Fig 1E). MAPK signaling emerged as one of the most significantly enriched pathways (Fig 1E), represented by several well-characterized KRAS effectors, including ARAF, BRAF, RAF1, PIK3CA, RASSF5, as well as GTPase activating proteins (GAPs) NF1, RASA1, RASA3, and receptor tyrosine kinases (RTKs) EGFR, ERBB2, and MET (Fig S1C). Beyond these canonical KRAS interactors, we identified several functionally relevant MAPK signaling proteins in close proximity to KRAS whose direct biophysical interactions with KRAS had not been reported (Fig S1C), including the key RAS activity regulator RASA3 and the RAS effector proteins PIK3CD. This finding suggests improvement in sensitivity of PPI screening techniques can enhance our understanding of even well-studied signaling pathways. In addition to MAPK signaling, we found several biological functions outside of signaling transduction as being significantly represented, including glucose metabolism, cell-cell communication, and membrane trafficking (Fig 1E). Detection of novel proteins within the well-characterized RAS signaling strengthened our confidence in uncovering novel KRAS interactors and underscored the utility of a highly sensitive and comprehensive protein interactome profiling approach.

### Dynamic change in KRAS proximal proteins in response to pharmacological RAS inhibition

The subset of 544 proteins whose proximity to KRAS susceptible to RASi was selected to analyze the changes in KRAS interactome following pharmacological perturbation (Fig 2A). Among these, 256 proteins (47.1%) exhibited significant decreases in KRAS proximity when treated with either panKRAS-off or panRAS-ON across KRAS mutation types, suggesting their interactions with KRAS were disrupted by RASi (Fig 2A). Conversely, 203 proteins (37.3%) displayed significantly increased proximity to inhibitor-bound KRAS (Fig 2A). Beyond these primary groups, 31 proteins were significantly downregulated by panRAS-ON but upregulated by panKRAS-off, while the opposite was observed for 10 proteins, suggesting potential protein interactions exceeding the binding to active GTP-bound KRAS.

**Fig. 2.**
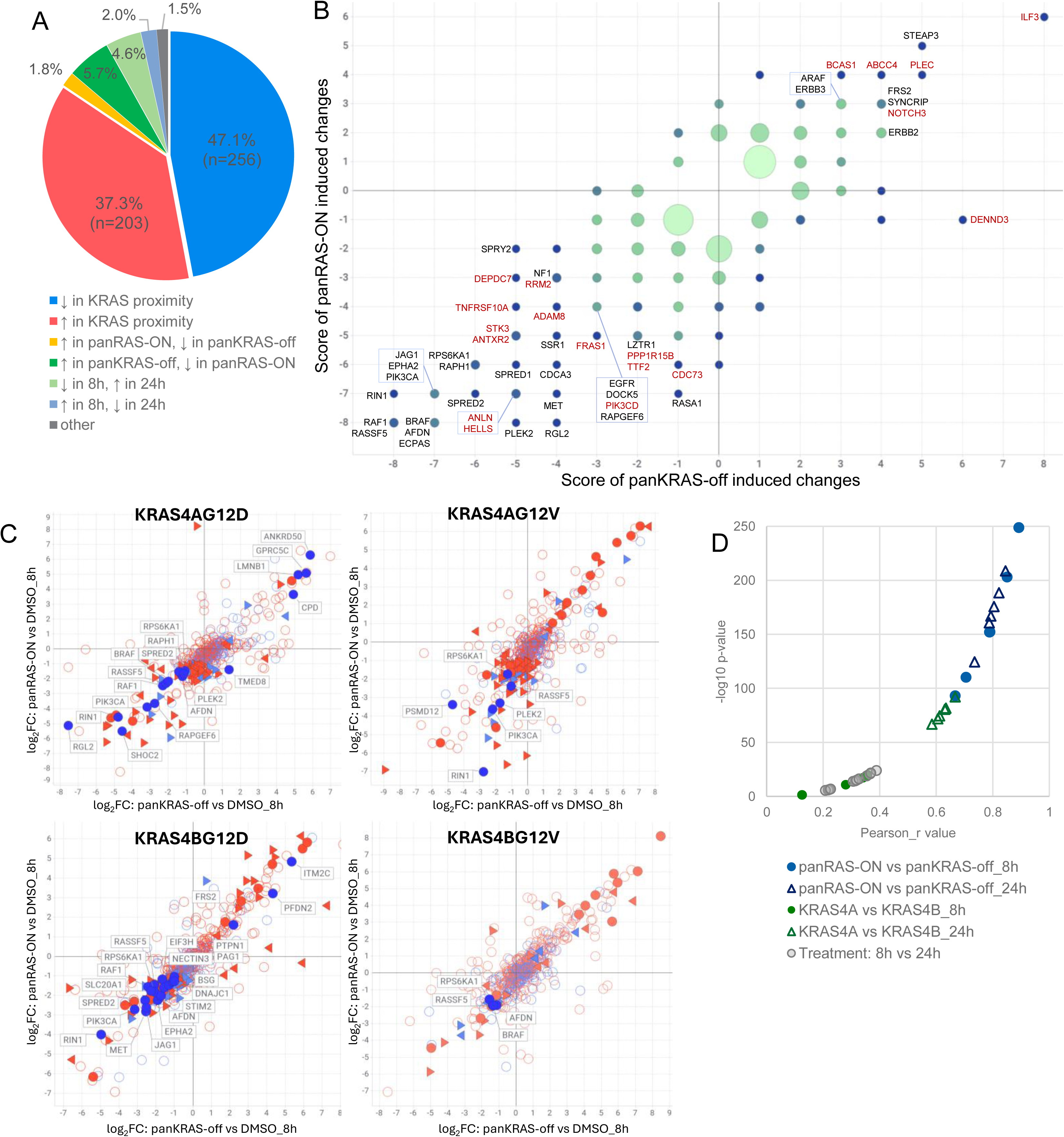
Remodeling of KRAS interactomes induced by RAS inhibitors (A) Composition of the subset of 544 RASi-sensitive KRAS proximal proteins grouped by abundance changes influenced by RASi treatment. (B) Dynamic KRAS interactome susceptible to pharmacological perturbation. Top 50 KRAS proximal proteins most significantly affected by RASi are labeled (black: proteins with published KRAS biophysical interactions; red: novel KRAS-proximal proteins first detected by our BioID study). Location of each protein in the coordinate system is determined by the designated scores based on the frequency of significant downregulation (negative score, log2FC<-1, FDR<0.05) or upregulation (positive score, log2FC>1, FDR < 0.05) detected in cells treated with either of the two RASi compared to corresponding DMSO-treated controls across the entire experiment. (C) Correlation of RASi-induced interactome changes between the two classes of RASi for oncogenic KRAS mutants. Log2FC of individual protein from cells expressing KRAS –G12D or –G12V mutants caused by panRAS-ON treatment for 8 hours were plotted against the log2FC caused by panKRAS-off in the same cells. Proteins significantly affected by both RASi with published biophysical KRAS interaction were annotated. Markers of individual proteins were categorized by published KRAS interaction and statistical significance of RASi-induced change. Blue: proteins with published KRAS interaction from BioGrid; red: novel KRAS proximal proteins identified by BioID; solid circle: proteins significantly affected by both RASi (FDR<0.05); triangle: proteins significantly affected by panRAS-ON (left pointed) or panKRAS-off (right pointed); hollow circle: proteins showed no statistical significance treated with RASi. (D) Correlation coefficient and statistical significance of RASi-induced log2FC from multiple Pearson correlation analysis of KRAS interactome regarding the type of RASi (blue), the type of KRAS splice variant (green), and treatment duration time (gray).

To further explore the frequency and direction of changes in interactions affected by RASi, we established a coordinate system for the dynamic KRAS interactome. Each KRAS-proximal protein was assigned to a negative or positive score based on the frequency of significant downregulation or upregulation (FDR < 0.05, log2FC > 1.0 or < -1.0) resulted from RASi treatment across the entire experiment. The position of each protein within the matrix is determined by its scores from the two RASi, providing a comprehensive view of how the KRAS interactome is reshaped by inhibitors functioning through distinct MOAs (Fig. 2B). The 256 proteins with decreased proximity to KRAS are localized in the third quadrant, while the 203 upregulated proteins are in the first quadrant. The top 50 most significant KRAS-proximal proteins were annotated (Fig. 2B), including well-characterized RAS effectors (e.g., RAF and PI3K proteins), regulators of KRAS activity such as GEFs and GAPs (e.g., NF1, RASA1, and RAPGEF6), and upstream RTKs (e.g., EGFR and MET). The reduced proximity of these proteins to KRAS in response to RASi reinforces the reliability of our findings in identifying robust and novel KRAS effectors.

The 203 proteins significantly upregulated in KRAS proximity upon RASi treatment include both classical KRAS interactors such as ARAF, ERBB2, and ERBB3 and potential novel KRAS interactors (ILF3, BCAS1, ABCC4, PLEC etc). To investigate the source of this upregulation, we examined the expression level of EGFR, ERBB2 and ERBB3 in H358 cells transduced with TurboID-KRAS4BWT or -KRAS4BG12V along with the parental H358 cells. The cells were treated with three RASi (panRAS-ON, panKRAS-off, KRASG12Ci sotorasib), panRAF inhibitor LXH254, and MEK inhibitor trametinib with treatment durations ranging from 30 minutes to 24 hours (Fig S2A). ERBB2 and ERBB3 were unchanged until 24 hours, when upregulation was observed in cells treated with RASi and MEKi (Fig S2A), suggesting the upregulation is not unique to KRAS inhibition. EGFR expression remained unaffected, suggesting reduced proximity of EGFR to KRAS detected by BioID reflects loss of interaction. Given that the significant RASi-induced upregulation of ERBB2 and ERBB3 in BioID was observed only in the 24-hour experiment (Fig S2B), we hypothesize that their increased proximity to KRAS is likely a result of compensatory feedback due to suppression of the MAPK pathway, rather than exclusively direct interactions with KRAS. Further analysis of the timing of upregulation showed that 67.5% of the RASi-induced increase occurred in the 24-hour experiment (137 of 203), compared to 30.1% in the RASi-induced decrease (77 of 256) (Fig S2C). These findings indicate that the short-term response of the KRAS interactome (8 hours) differs from the longer-term response observed at 24 hours. This distinction highlights the dynamic and temporal nature of the KRAS interactome’s adaptation to pharmacological inhibition.

### LMW RAS inhibitors with distinct MOA affect the same repertoire of KRAS PPIs

The distribution of KRAS interactors within the coordinate system of our KRAS interactome indicated a potential positive correlation in KRAS interactome change in response to the two classes of RASi (Fig 2B). Plotting the fold-changes of individual KRAS-proximal proteins affected by panRAS-ON versus panKRAS-off revealed significant positive correlations in the disrupted protein interactions between distinct MOAs, with canonical KRAS interactors being particularly prominent (Fig. 2C). This correlation was consistently observed across all tested KRAS variants (Fig S3A), suggesting the KRAS interactors blocked by CYPA at the binding interface of active GTP-bound KRAS substantially overlap with those dissociated due to the accumulation of inactive GDP-bound KRAS. On the contrary, the correlation between 8- and 24-hour treatment within a given condition was comparatively weaker (Fig. S3B), as were correlations between the interactomes of KRAS4A and KRAS4B (Fig. S3C). Interestingly, both the statistical significance and the correlation coefficient between the two splice variants (KRAS4A and KRAS4B) were notably higher at the 24 hour compared to the 8 hour time point (Fig. 2D). To better understand the differences, we categorized the 544 KRAS-proximal proteins based on their interaction with KRAS4A or KRAS4B. Despite the high degree of sequence and functional redundancy between the two isoforms, only 16.7% of RASi-sensitive KRAS interactors were shared between KRAS4A and KRAS4B after 8 hours of RASi treatment (Fig S3D). The percentage of overlap increased at the 24 hour time point for both downregulated and upregulated proteins, suggesting that the feedback response to KRAS suppression for the two splice variants is similar. Consistent with the higher correlation observed between the KRAS4A and KRAS4B interactomes at 24 hours (Fig 2D), we conclude that the distinct protein networks recruited by KRAS4A and KRAS4B exhibit similar responses to long-term pharmacological inhibition of KRAS, regardless of the modality.

### AFDN associates with oncogenic KRAS mutants

AFDN was a top KRAS interactor across wild-type and mutants of both KRAS4A and KRAS4B in our BioID study (Fig 2B). The degree of AFDN displacement from KRAS proximity by RASi was comparably significant to that of canonical KRAS effectors such as BRAF and CRAF (Fig 3A). AFDN is a multidomain scaffold protein crucial for the formation and maintenance of adherens junctions and tight junctions, achieved through interacting with various binding partners such as nectin and the E-cadherin adhesion systems (17–19). *Afdn* knockout mice exhibit embryonic lethality due to developmental defects, underscoring the indispensable role of AFDN in regulating cell polarity and cell-cell adhesion (19,20). First identified in 1995 when it was co-purified with HRAS from bovine brain extracts, AFDN has been characterized as a distinct effector protein for the RAS family, directly binding to HRAS, RAP1A and RAP2A (21,22). More recently, proximity labeling-based screening techniques have provided additional evidence linking AFDN to KRAS (8,23).

**Fig 3.**
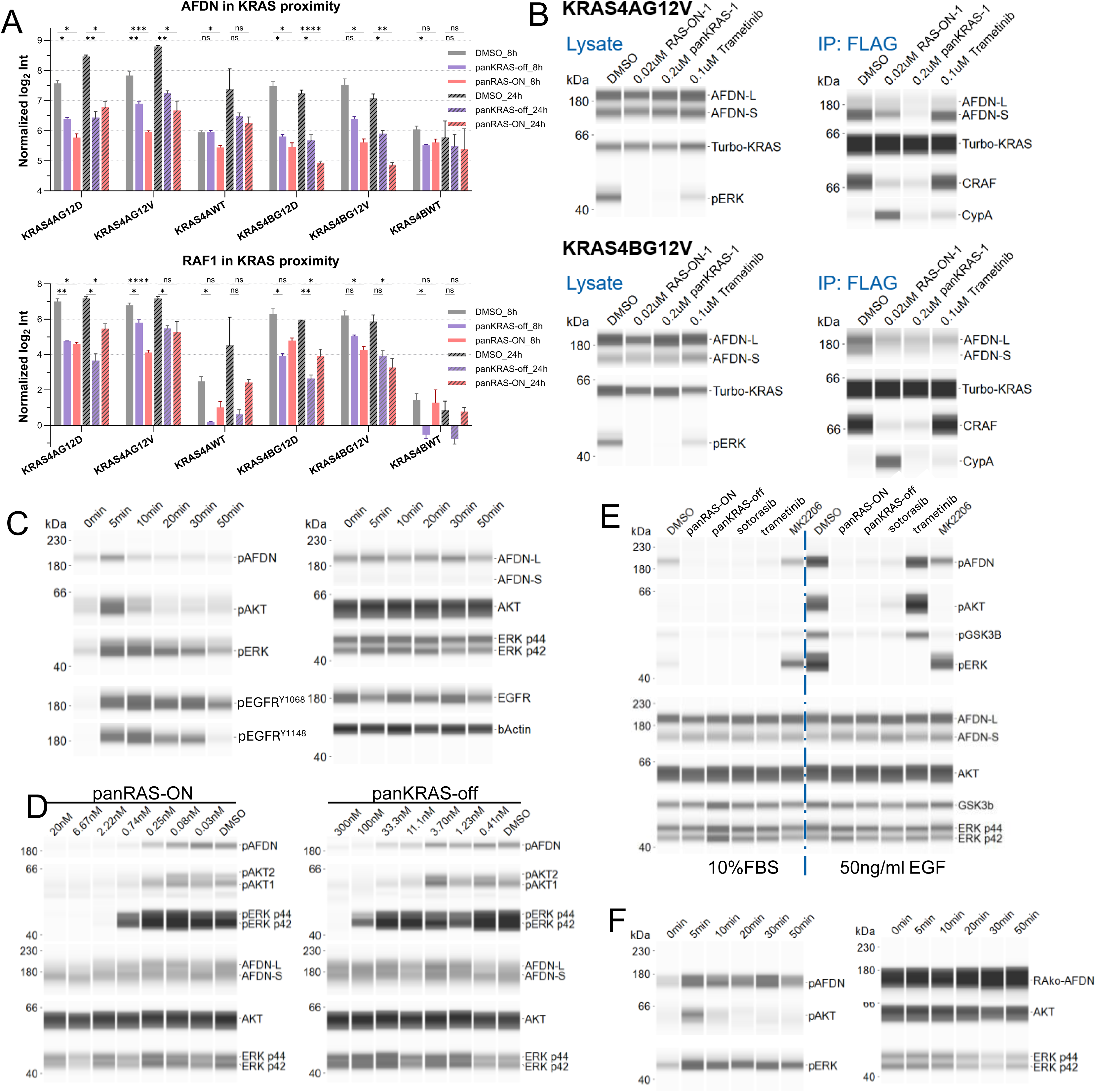
KRAS interacts with AFDN and regulates EGF-stimulated phosphorylation of Ser1718 of AFDN **(A)** Abundance of AFDN and RAF1 in KRAS proximity were significantly decreased by both RASi in BioID. (✱: FDR<0.05; ✱✱: FDR<0.005; ✱✱✱: FDR<0.0005; ✱✱✱✱: FDR<0.0001) **(B)** Immunoblot detection of KRAS with co-immunoprecipated AFDN and RAF1 from H358 cells expressing FLAG-tagged TurboID-KRASG12V of either KRAS4A or KRAS4B treated with two RASi and trametinib. **(C)** Immunoblot of pAFDN^S1718^ stimulated by 100 ng/ml EGF in H358 cells. p-ERK, p-AKT and p-EGFR were blotted to show the stimulatory effect of EGF on MAPK pathway. **(D)** Immunoblot of pAFDN^S1718^ suppressed by panRAS-ON and panKRAS-off in a dose-dependent manner in H358 cells. **(E)** EGF-activated pAFDN^S1718^ in H358 is completely blocked by three RASi (0.02 µM panRAS-ON, 0.1 µM panKRAS-off, 0.1 µM KRASG12Ci) but less effective by trametinib (0.1 µM) and AKTi (MK2206, 1 µM). **(F) I**mmunoblot of pAFDN^S1718^ fragment (aa350-aa1824) in shRNA-induced AFDN knockdown cells. Cell lysates were collected from cells treated with 100 ng/ml EGF at 48 hours post-transient transfection of AFDN (350-1824)

Our results demonstrate that KRAS^G12D^ or KRAS^G12V^ -transduced H358 cells exhibit a higher baseline abundance of AFDN in the proximity of KRAS compared to wild-type KRAS-expressing cells, suggesting that AFDN may have a higher affinity for GTP-bound active KRAS than GDP-bound inactive KRAS (23). To independently confirm the direct protein interaction between AFDN and KRAS, the FLAG-tagged TurboID-KRAS^G12V^ fusion proteins expressed from H358 cells were immunoprecipitated with the anti-FLAG antibody. AFDN was co-immunoprecipitated with ectopic KRAS^G12V^ but not in cells treated with the inhibitor panRAS-ON or panKRAS-off (Fig 3B), suggesting the protein interaction between KRAS and AFDN is sensitive to both RAS inhibitors.

### KRAS mediates EGF-stimulated phosphorylation of AFDN

Enrichment analysis of the RASi-sensitive KRAS interactome identified the cell-cell communication pathway as significantly enriched (Fig. 1E). Mapping the RASi-susceptible KRAS interactome with components of adherens junction and tight junction showed a substantial overlap with both protein complexes (Fig S4). The entire scaffold signaling complex recruited by junctional adhesion molecule-A (JAM-A) were biotinylated, comprising ZO-2, AFDN, and PDZGEF1/2, of which the latter two were significantly changed in KRAS proximity upon RASi treatment (Fig. S4), with similar impact on the nectin-recruited adherens junction complex. The overlap between KRAS interactors and the AFDN interactome identified by an independent study further supports the hypothesis that KRAS participates in regulating tight junctions through AFDN (Fig. S5) (23). However, the specific mechanism by which the interaction between AFDN and KRAS contributes to its biological function remains unclear.

AFDN has multiple phosphorylation sites that modulate its interactions with binding partners and influence cellular processes, such as migration (24,25), adhesion (26), and signaling (27,28). The most studied AFDN phosphorylation site is Ser1718 (24,27,29), located in the actin-binding domain. This site conforms to the optimal AKT consensus substrate motif (RXRXXS/T) (24,30). Furthermore, the sequence surrounding Ser1718 is evolutionarily conserved in AFDN from Drosophila to mammals (24), highlighting its importance as a key regulatory feature in AFDN’s biological activity.

Given that the PI3K/AKT pathway is a key component of the KRAS-regulated signaling network, we tested whether phosphorylation at this site could be activated by EGF stimulation in H358 cells. Serum-starved H358 cells treated with 50 ng/mL EGF exhibited significant induction of a series of RAS/MAPK pathway-related phosphorylation events, including p-EGFR (Y1068 and Y1148), p-ERK, p-AKT, and notably p-S1718 of AFDN (Fig 3C). Phosphorylation of AFDN^S1718^ peaked concurrently with p-AKT at 5 minutes post-EGF stimulation, while p-ERK peaked slightly later at 10 minutes with a longer duration of activation. The activation of p-AFDN^S1718^ by EGF prompted further investigation into the role of KRAS in this process. We evaluated the impact of KRAS on modulating AFDN phosphorylation by treating the H358 cells with the two RAS inhibitors. Both RASi suppressed p-AFDN^S1718^ in a dose-dependent manner although panRAS-ON is more effective than panKRAS-off, probably due to the difference in potency (Fig 3D). Both p-AFDN and p-AKT were completely suppressed while p-ERK was only partially suppressed at the same concentration of RAS inhibitor, suggesting that p-AFDN is more likely to be regulated by AKT rather than downstream of ERK activity.

To determine whether this suppression of phosphorylation resulted from direct inhibition of KRAS activity or from downstream effects of the MAPK pathway, we treated H358 cells with a panel of inhibitors targeting KRAS (panRAS-ON, panKRAS-OFF, sotorasib), MEK (trametinib), and AKT (MK2206) (Fig 3E). Serum-starved H358 cells were treated with inhibitors for 1 hour, followed by stimulation of 10% FBS or 50 ng/ml EGF for 5 minutes. EGF was substantially more efficient at activating the MAPK pathway in H358 cells than 10% FBS as shown by the markedly higher levels of p-ERK, p-AKT, p-GSK3b and p-AFDN than in the control group (Fig 3E). All three RASi completely blocked the phosphorylation of AFDN^S1718^ as well as p-ERK and p-AKT in H358 cells. In contrast, the pan-AKT inhibitor MK2206 only partially reduced the level of p-AFDN^S1718^. Trametinib had no effect other than suppressing p-ERK as the direct substrate of MEK kinases. The degrees of pathway inhibition (pERK in trametinib-treated cells, pAKT/pGSK3B in MK2206-treated cells) confirmed that the incomplete suppression of p-AFDN by MEK and AKT inhibitors cells was not due to inadequate inhibition but rather suggested that other kinases were partially responsible for phosphorylation of Ser1718.

The fact that only RAS inhibitors fully suppressed p-AFDN^S1718^ indicates that KRAS is the primary regulator of this phosphorylation event. Furthermore, a global phosphoproteomic study by Klomp *et al*. revealed significant downregulation of p-AFDN^S1718^ in the KRAS^G12C^ mutant lung cancer cell line H358 and colorectal cancer cell line SW837 when treated with the KRAS^G12C^ inhibitor MRTX1257 (Fig S6A) (31). Previous studies with sequential inhibition of three oncogenic RTKs (c-Met, EGFR, and PDGFRα) and their downstream serine-threonine kinases (PI3K, MEK and MTOR) with clinical-relevant inhibitors showed that p-AFDN^S1718^ is substantially reduced by RTK inhibitors (Gefitinib, Gleevec, and SU11274), partially decreased by the PI3K inhibitor wortmannin, and unaffected by MEK (U0126) or MTOR (rapamycin) inhibitors (32), further endorsing the primary regulatory role of KRAS on AFDN phosphorylation. Together, these findings support the conclusion that KRAS acts as a primary upstream regulator of AFDN^S1718^ phosphorylation through EGFR signaling, likely by recruiting a range of kinases, including but not limited to AKT.

Following our co-IP confirming the interaction between AFDN and KRAS, we sought to determine whether KRAS-dependent phosphorylation of AFDN^S1718^ requires direct KRAS-AFDN binding. A deletion variant lacking the N-terminal RA1/2 domain, termed RAΔAFDN (aa350-1824), was expressed in H358 cells in which endogenous AFDN had been silenced through RNA interference (RNAi; shAFDN1-3) (Fig. S6B). Immunoblot confirmed successful expression of the RAΔAFDN protein in the *AFDN*-knockdown cell line. Upon stimulation with 100 ng/ml EGF, rescued cells expressing RAΔAFDN exhibited phosphorylation of AFDN^S1718^ to a similar degree as cells expressing wild-type AFDN (Fig. 3F), indicating the RA1/2 domain was not required for this phosphorylation event.

### AFDN-dependent directional cell migration

RAS-driven cancers are often characterized by a propensity for metastasis and malignancy, with key steps in invasion and metastasis involving changes in cell adhesion and enhanced motility (33). As a critical regulator of cell polarity and adhesion, AFDN has been linked to cell migration, invasion and metastasis in several cancer types (34,35). To investigate the impact of the oncogenic mutant KRAS^G12C^ on cell motility, we tracked H358 cells treated with the two RASi over a 48 hour period. Cells were seeded at low-density to allow free movement of individual cells. Both RASi reduced the speed of individual cell movement with complete suppression observed at 0.74 nM panRAS-ON and 300 nM panKRAS-off (Fig 4A). Notably, cell motility was effectively suppressed before MAPK signaling was fully blocked, shown by the residual p-ERK in cells treated with 0.74 nM panRAS-ON (Fig S7A). Furthermore, treatment with 0.74 nM panRAS-ON resulted in over 90% of the nuclei remaining healthy, in contrast to the significant decrease in cell viability seen at the highest dose of 20 nM (Fig S7B). This illustrates that low concentrations of panRAS-ON can halt cell division, reducing the total cell population by 60% without inducing substantial toxicity (Fig S7C). Similarly, H358 cells retained >85% healthy nuclei even when treated with 300 nM panKRAS-off (Fig S7C), and individual cell motility was fully suppressed at 1.23 nM of panKRAS-off (Fig 4A) despite active MAPK signaling (Fig. S7A). These findings suggest that pharmacological inhibition of KRAS can impair cell motility without inducing cell death.

**Fig 4.**
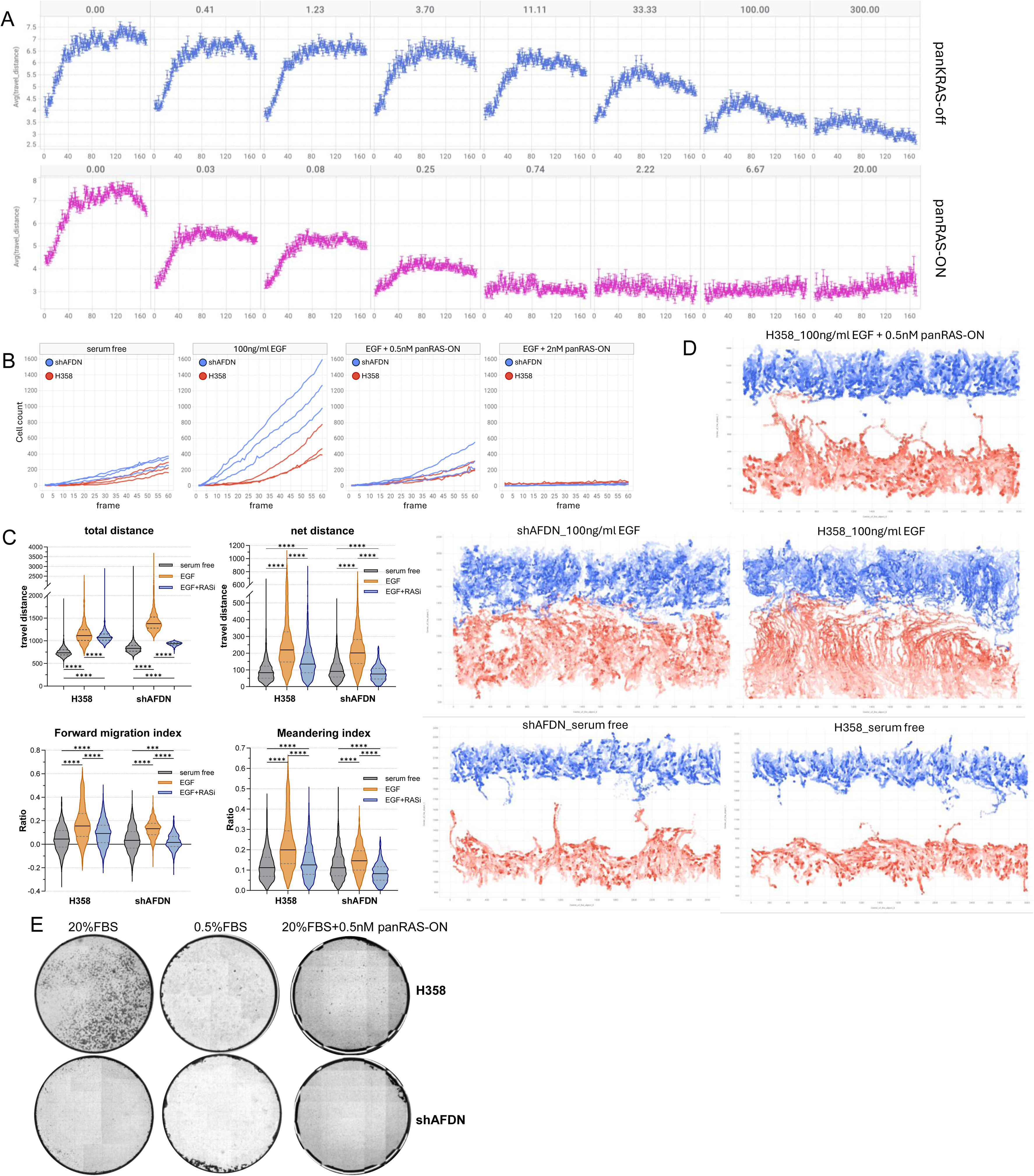
KRAS coordinates with AFDN to regulate the EGF-activated collective migration **(A)** RAS inhibitors suppressed the speed of cell motility in free movement. Average speed of individual cells per well was calculated from H358 cells treated with panRAS-ON (starting at 20 nM, bottom) or panKRAS-off (starting at 300 nM, top) at different concentrations in triplicate. **(B)** Number of cells entering the wound area during the first 16 hours of the wound healing process between H358 cells and the shAFDN-knockdown cells under different conditions. Both cell lines were grown in serum-free media during the wound healing process supplemented with 100 ng/ml EGF or additional panRAS-ON (0.5 nM or 2 nM) in triplicate. **(C)** Track measures calculated from aggregated individual cell tracking. Parameters were calculated using equations from Svensson et al (16). FMI (forward migration index): ratio of the progress in the upright direction by a cell to the total distance. Meandering index: ratio of net distance to total distance per individual cell. Statistical significance was analyzed by one-way ANOVA with Tukey correction for multiple comparisons by GraphPad Prism. (✱: FDR<0.05; ✱✱: FDR<0.005; ✱✱✱: FDR<0.0005; ✱✱✱✱: FDR<0.0001) **(D)** Representative migratory pattern of individual cell tracking during the wound healing process. The migratory track of 500 cells (250 on each side of the wound per condition) trackable across the entire 48 hour experiment were selected to reflect the migration pattern. **(E)** Transwell migration assay of H358 and shAFDN-knockdown cell lines. Both cell lines were seeded in DMEM media supplemented with 0.5% FBS. Media containing 20% FBS with or without RASi were used as chemoattractant.

To determine whether KRAS-mediated motility in H358 cells depends on AFDN, we evaluated the effect of *AFDN* loss on collective cell migration using a wound-healing assay with *AFDN* knockdown (KD) and parental H358 cells. In serum-free media, we monitored wound closure in response to EGF over a 48-hour time period. Silencing *AFDN* in H358 cells did not impair the ability of cells to respond to EGF stimulation and migrate from the overconfluent area to the wound. Both parental H358 and *AFDN* KD cells completely closed the wound after 48 hours when treated with 100 ng/ml EGF (Supplementary video, Fig 3C). Additionally, the stimulatory effect of EGF on cell migration was fully inhibited by 0.5 nM panRAS-ON in both cell lines, indicating that EGF-driven collective cell migration is mediated by KRAS but does not require AFDN.

Although both cell lines achieved complete wound closure under EGF stimulation, the specific mechanisms acting in each cell line differed, as revealed by our in-depth quantitative analysis. During the first 16 hours of wound healing, the number of migratory cells entering the wound under EGF stimulation was significantly higher for *AFDN* KD cells compared to parental H358 cells (Fig 4B), although both cell lines reached a similar peak cell density by 48 hours (Fig S8A), suggesting wild type cells required more time to initiate collective migration of the wound-healing process. Moreover, the total distance of individual cells migrating over the 48 hours of the wound-healing process for EGF-activated AFDN KD cells was significantly higher than H358 cells in the same condition (Fig 4C). In contrast, no significant differences in cell numbers were observed between the two cell lines under serum-starvation or RASi-treated conditions. These findings suggest that RNAi-induced silencing of AFDN promoted cell motility in response to EGF stimulation compared to the parental H358 cells.

Detailed tracking analysis of individual cells revealed that H358 cells close the wound with greater directionality compared to AFDN depleted cells. The migratory track of individual cells explicitly demonstrated that EGF stimulation enhanced both the migration distance and linearity of H358 cells during the wound healing process (Fig 4D). In contrast, AFDN depleted cells closed the wound through movement resembling Brownian motion rather than directed migration under EGF stimulation (Fig 4D), suggesting AFDN is not required for cell motility but is essential for regulating migratory directionality. Although AFDN KD cells traveled a higher total distance upon EGF stimulation, the net and max distance in H358 cells were significantly higher; the forward migration index and meandering index showing a similar trend (Fig 4C, Fig S8B). The lower turning angle observed in H358 cells compared to AFDN KD cells during migration further showed the lack of orientation in AFDN KD cells (Fig S8B). This suggests that H358 cells maintain sustained polarity in their movement over time, whereas this polarity is lost in AFDN KD cells.

Consistent with previous findings, RASi significantly suppressed the collective migration, as neither cell line was able to close the wound when treated with 0.5 nM panRAS-ON or higher concentration (Fig 4D, Supplementary video), suggesting the EGF-activated enhancement of directional migration requires both functional KRAS and AFDN. The motility and directionality of H358 cells were inhibited to baseline levels comparable to those observed under serum-starvation conditions (Fig 4C). The inhibitory impact of panRAS-ON was even greater on *AFDN* KD cells, as we found the trackability of individual cells significantly dropped whereas the number of segmented cells per frame remained comparable to counterparts (Fig S9A, S9B), likely due to overcrowded cell density resulting from deprived motility by RASi. These findings indicate that EGF-stimulated collective migration is mediated by KRAS in lung cancer cell lines harboring oncogenic KRAS mutations. The dependency of H358 cell directionality on AFDN was further confirmed using a transwell assay, where depletion of AFDN completely abolished the chemotactic migration observed in parental H358 cells (Fig 4E, Fig S10). Collectively, these findings highlight AFDN as a critical mediator of cell migration directionality and polarity, playing a key role in enabling cells to respond to spatial and biochemical stimuli effectively.

## Discussion

Since BioID was first applied to KRAS in 2018 (9), the KRAS regulatory network has broadened with new interactors identified, including mTORC2 (4), PIP5K1 (9), RSK1 (36), and BIRC6 (37). Efficient protein interaction screening by BioID enabled researchers to discover novel interactors and compare networks associated with wild-type versus mutant KRAS (38), as well as distinct RAS paralogs (39). Early BioID studies used the first-generation biotin ligase BirA* in common tool cell lines (HEK-HT, HEK293) (9,39). Beyond the dependency of BirA* on high biotin concentration and long labeling times, these cell lines do not reflect KRAS-driven cancer contexts. We employed TurboID, the latest engineered BirA* with markedly higher activity (12,40), in a KRAS^G12C^ non-small cell lung cancer model to establish a context-relevant KRAS interactome. Using a 3-hour biotin pulse, we stably detected more than 2,000 proteins proximal to each KRAS variant.

KRAS was considered "undruggable" until the development of covalent KRAS^G12C^ inhibitors targeting the Switch II pocket. More recently, noncovalent Switch II binders have been designed, such as the G12D-selective inhibitor MRTX1133 (41) and panKRAS inhibitors like Pan KRas-IN-1. Binding to the Switch II pocket induces conformational changes in KRAS’s Switch I and II loops, which are critical for recruiting downstream effectors (41). For instance, MRTX1133 disrupts the KRAS(G12D)-GTP-RAF1 complex, inhibiting MAPK pathway activation. Conversely, RMC-6236 inhibits KRAS-GTP by forming a ternary complex with RAS and Cyclophilin A, disrupting KRAS-CRAF complexes in cells.

Using two mechanistically distinct RASi, we characterized, for the first time, the dynamic KRAS interactome susceptible to small-molecule perturbation and the associated molecular changes. We observed a strong positive correlation in inhibitor-induced interactome shifts between panRAS-ON and panKRAS-off across all KRAS variants, indicating that distinct MOA targeting different nucleotide states of KRAS disrupt similar interactions. The immediate impact of panKRAS-off was smaller than that of panRAS-ON, but their longer-term effects converged, suggesting that trapping KRAS in its inactive state via a switch II pocket binder can be functionally equivalent to forming a ternary complex between GTP-bound KRAS and CypA.

Chemoproteomic profiling of inhibitor-induced changes not only revealed KRAS interactors sensitive to pharmacological perturbation but also highlighted inhibitor-insensitive cohorts. We identified numerous established KRAS interactors, including key regulators of KRAS activity and MAPK signaling, yet many were unexpectedly unaffected by RAS inhibition such as RALGDS, SOS1, and SHP2 (Fig S1C). Both GEFs and GAPs were robustly biotinylated; however, NF1 and RASA3 were displaced from KRAS proximity by RAS inhibitors, while GEFs remained largely unaffected. These findings suggest variable sensitivity among KRAS interactors to KRAS inhibition, likely reflecting functional differences. Additionally, 273 proteins were significantly upregulated near inhibitor-bound KRAS. Measuring ERBB2 and ERBB3 expression in H358 cells treated with RASi versus MEKi over 24 hours demonstrated that increased KRAS proximal biotinylation could be driven by compensatory upregulation following pathway suppression. These results underscore that proximity labeling may not exclusively report direct binding but can also reflect transcriptional, translational or stability changes in proteins.

AFDN emerged as the top RASi responsive protein, with statistical significance comparable to BRAF and RAF1. AFDN abundance was greater with KRAS^G12D^ and KRAS^G12V^ than with wild-type KRAS, suggesting preferential association with active GTP-bound KRAS over GDP-bound KRAS. Further probing revealed that phosphorylation of AFDN^S1718^, reported essential for promoting migration in T47D cells (24), is activated by EGF via KRAS signaling. Although Ser1718 is a direct AKT substrate, PI3K or AKT inhibition only partially reduced its phosphorylation in prior studies (24,32). In contrast, inhibiting KRAS, either with a pan-RAS inhibitor or with adagrasib specifically targeting KRAS^G12C^, fully blocked EGF-induced Ser1718 phosphorylation. These results place KRAS upstream of AKT as a central regulator of EGF-activated p-AFDN^S1718^ and imply additional Ser/Thr kinases participating independently of AKT.

The role of AFDN in cancer cell migration and metastasis have garnered increasing attention and appeared to be context-dependent (34). Our literature survey indicates that AFDN’s multifunctionality, context dependence, and variable definitions of “migration” across assays together better explain the apparent contradictions. Different experimental approaches, or even the same assay under varying conditions, interrogate distinct facets of migration and therefore capture only parts of AFDN’s functional complexity, yielding seemingly conflicting outcomes. For example, *AFDN* loss markedly inhibited migration in BT549 and MDA-MB-468 cells (24) but increased migration velocity in MCF7, SK-BR-3, and MDA-MB-231 cells (42), despite all being breast cancer lines. The observed differences were attributable to chemoattractant-driven transwell assays in the former versus wound-healing assays in complete media (10% FCS) in the latter. A more striking case is T47D, which naturally expresses low AFDN due to genetic alterations: AFDN overexpression enhanced migration in transwell assays (24) but significantly reduced both spontaneous and heregulin-β1-induced collective migration in wound healing (42).

Notably, the inhibitory effect of AFDN depletion in transwell assays was completely reversed in Matrigel invasion assays. AFDN knockdown increased invasiveness across several cancer types, including breast (SK-BR-3) (42), endometrial (Ishikawa, HEC1A) (43), osteosarcoma (U2OS) (44), and pancreatic (BxPC-3, HPAC) (20). Moreover, the pro-invasive effect of AFDN depletion in endometrial cells disappeared when assessed using transwells without Matrigel (43), indicating whether AFDN promotes or inhibits migration depends more on experimental conditions than on cancer type, reflecting AFDN’s intrinsic functional complexity.

We disentangled directionality from motility of cell migration by analyzing tracks of individual cells stimulated exclusively with EGF. Our high-resolution wound-healing analysis showed that AFDN contributes to orienting cells toward biochemical and biophysical cues. Motility was separable from directionality: shAFDN-knockdown cells, though aimless, completely closed the wound in response to EGF with more cells participating in migration at higher speed than wild-type cells. RASi completely abolished the EGF-induced motility in AFDN knockdown cells. Overall, we conclude that KRAS is a central regulator of EGF-activated directional migration, with AFDN mediating directionality downstream of KRAS signaling.

AFDN has been proposed as a suppressor of RAS-ERK signaling in several cancer cell lines with wild-type KRAS including MCF7 (23), SK-BR-3 (42), Ishikawa (43), U2OS (44), and NIH3T3 cells (28). In our KRAS-mutant H358 model, we observed no differences in the duration or magnitude of EGF-induced ERK phosphorylation between parental and shAFDN knockdown cells (data not shown), suggesting that AFDN may modulate RAS–ERK signaling in wild-type contexts but not in KRAS-mutant cancer cells, where MAPK signaling is constitutively active.

BioID can reveal changes in protein interactions proximal to highly dynamic signaling proteins, such as RAS, in relevant cancer contexts. A clearer understanding of small-molecule-induced interactome remodeling—particularly for drug targets like KRAS—can inform drug development and highlight candidates for synthetic lethality. Our study provides a comprehensive map of KRAS interactome dynamics under pharmacological perturbation. Future work should examine long-term rewiring under sustained KRAS inhibition to help explain and potentially overcome limited response rates and acquired resistance in current KRAS-targeted therapies.

## Supporting information

H358_serum free

shAFDN_serum free

H358+EGF

shAFDN+EGF

H358+EGF+RASi

shAFDN+EGF+RASi

---

**Supplementary Fig 1.**
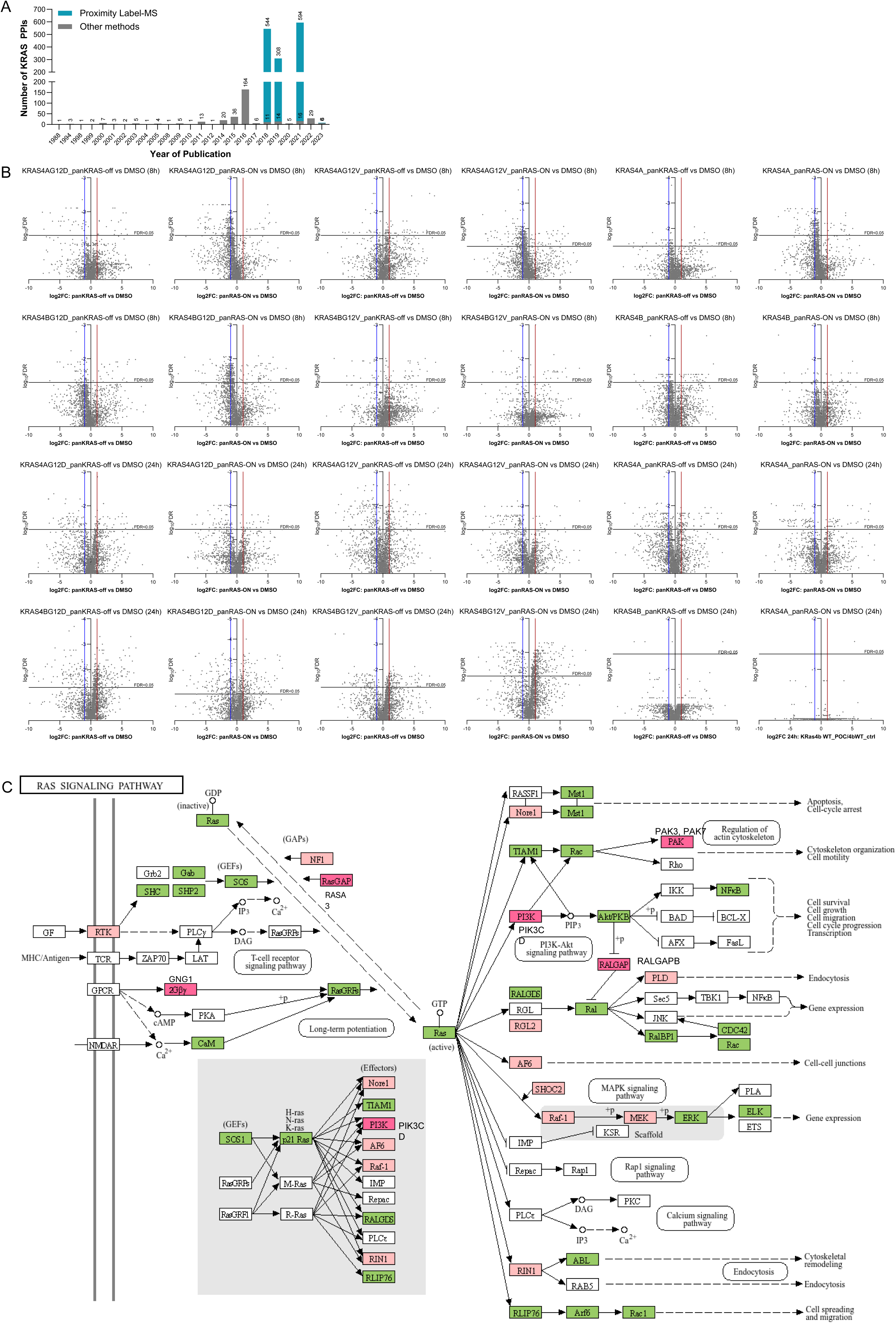
(A) Number of published KRAS biophysical interactions from BioGrid by the year of the publication. Number of protein interactions identified by Proximity Label-MS (teal) versus other methods (gray) were labeled accordingly. (B) Volcano plot of KRAS-proximal proteins in cells following 8 hours or 24 hours of RAS inhibition by panKRAS-off or panRAS-ON compared to corresponding DMSO-treated control across KRAS variants. Significant changes in the volcano plots were calculated using Student’s two-sided t-test, and the false discovery rate (FDR)-adjusted P-values were calculated using the Benjamini–Hochberg method. (C) Components of RAS signaling pathway from KEGG database. Proteins detected in BioID were labeled accordingly (green: biotinylated proteins; light pink: KRAS proximal proteins significantly affected by RASi; dark pink: RASi-susceptible KRAS interactors whose biophysical proximity with KRAS were first observed in our BioID study, including RASA3, GNG12, PIK3CD, PAK3, PAK7, RALGAPB).

**Supplementary Fig 2.**
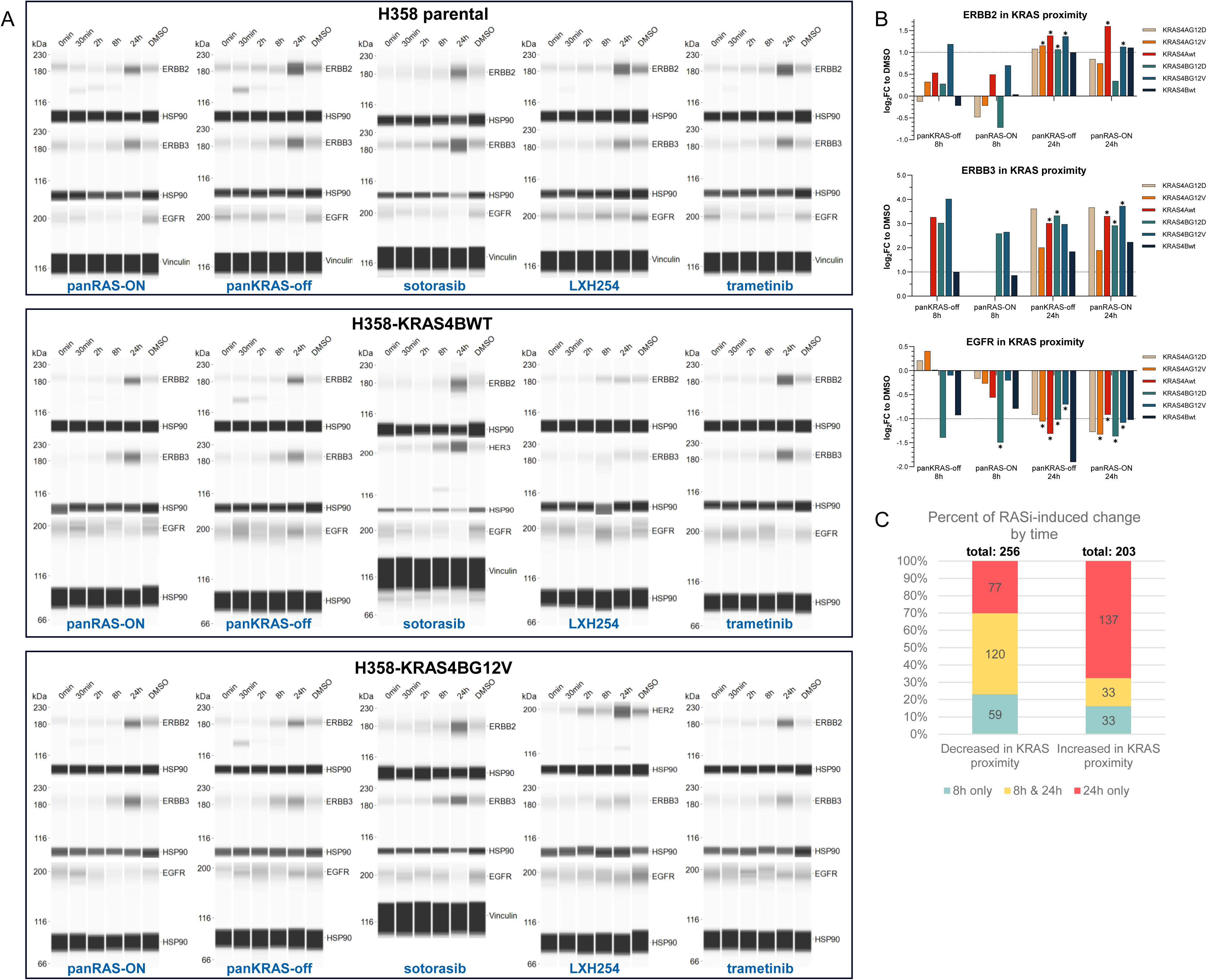
(A) Immunoblot for ERBB2, ERBB3 and EGFR in H358 cells ectopically expressing wild-type KRAS4B, KRAS4B^G12V^ or the parental H358 cells treated with small molecule inhibitors targeting essential components of the MAPK pathway. (B) Log2FC of ERBB2, ERBB3 and EGFR in KRAS proximity in response to RASi in H358 cells expressing TurboID-fused KRAS variants identified by BioID. (✱: FDR<0.05) (C) Percentage of RASi-induced downregulation and upregulation in KRAS interactome categorized by time of experiment. Number of significant differences occurred exclusively in the 8-hour experiment, 24-hour experiment, or both RASi-treatment were listed.

**Supplementary Fig 3.**
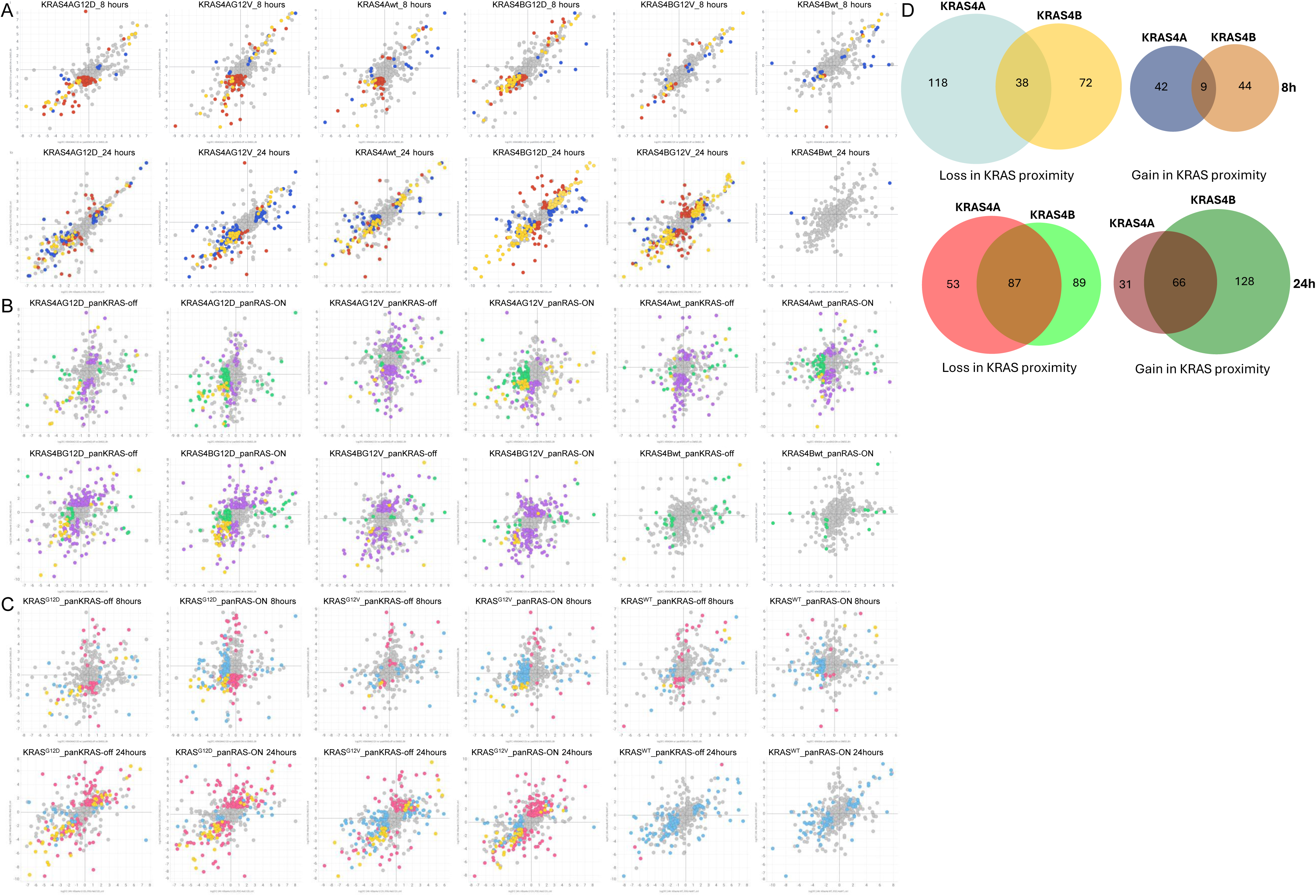
(A) Log2FC of individual KRAS-proximal protein in H358 cells treated with panRAS-ON compared to corresponding DMSO-treated control group plotted against the Log2FC for panKRAS-off for each KRAS variant in the BioID experiment. Proteins were colored based on the statistical significance of each fold-change: yellow: proteins significantly affected by both panRAS-ON and panKRAS-off (FDR<0.05, log2FC>1 or <-1); red: proteins significantly affected by panRAS-ON only; blue: proteins significantly affected by panKRAS-off only. (B) log2FC of individual KRAS-proximal protein affected by RASi for 8 hours were plotted against the changes affected by the same RASi at 24 hours. Yellow: proteins significantly affected by the same RASi at both 8 hour and 24 hour experiments. Green: proteins significantly affected by the RASi at 8 hours only. Purple: proteins significantly affected by the RASi at 24 hours only. (C) log2FC of individual KRAS-proximal protein affected by RASi in KRAS4A were plotted against the change occurred under the same condition and the same mutant in KRAS4B. yellow: proteins significantly affected by RASi at the proximity of both KRAS4A and KRAS4B; blue: proteins significantly affected by RASi at the proximity of KRAS4A only; pink: proteins significantly affected by RASi at the proximity of KRAS4B only. (D) Venn diagram of the numbers of shared and unique RASi-sensitive KRAS proximal proteins of KRAS4A and KRAS4B at 8 hour and 24 hour experiments, respectively.

**Supplementary Fig 4.**
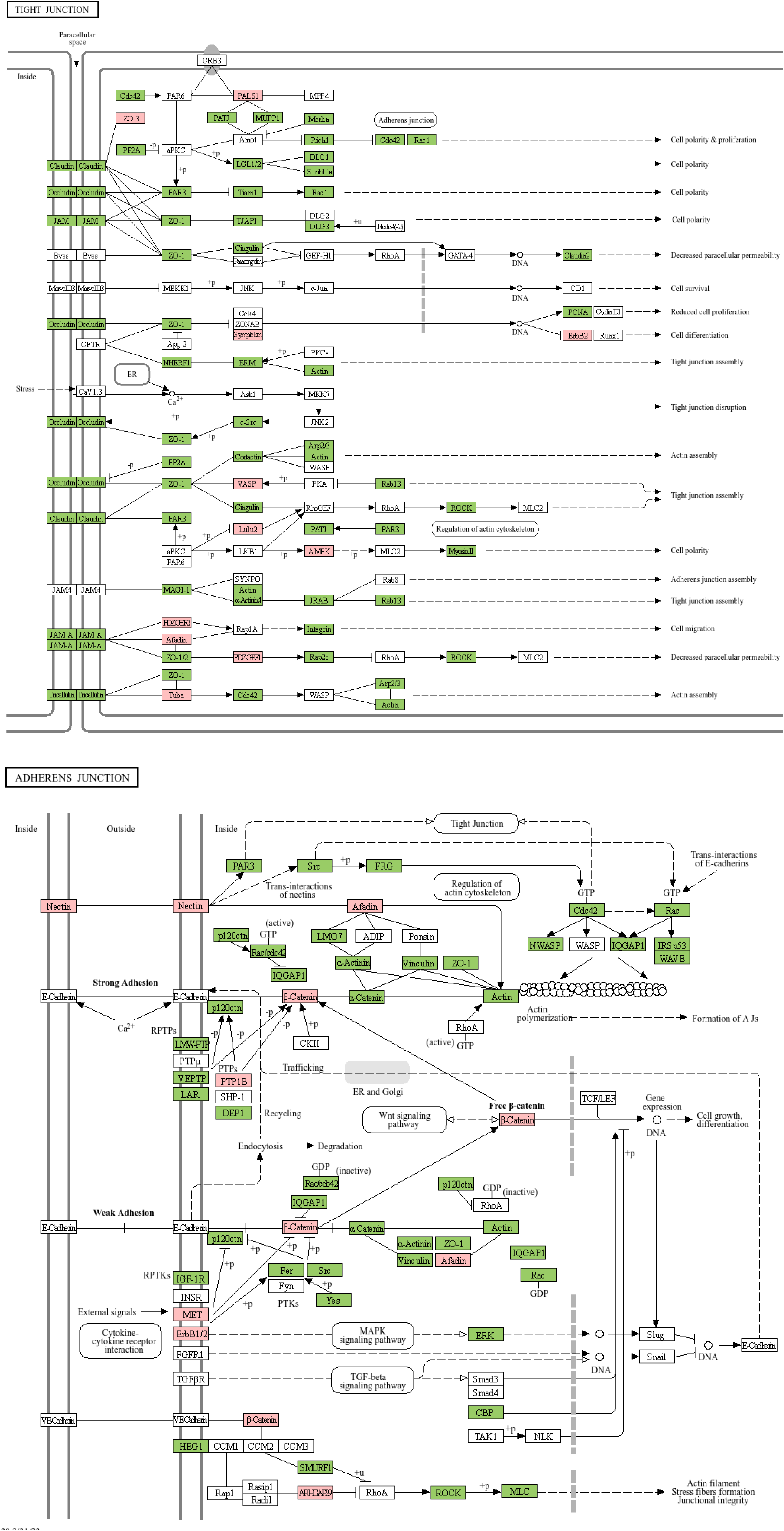
Overlap of BioID-identified KRAS interactome with the adheren junctions and tight junctions pathway from KEGG database. Proteins detected in BioID were labeled (red: biotinylated proteins; pink: biotinylated proteins significantly affected by RASi in the KRAS proximity, FDR<0.05).

**Supplementary Fig 5.**
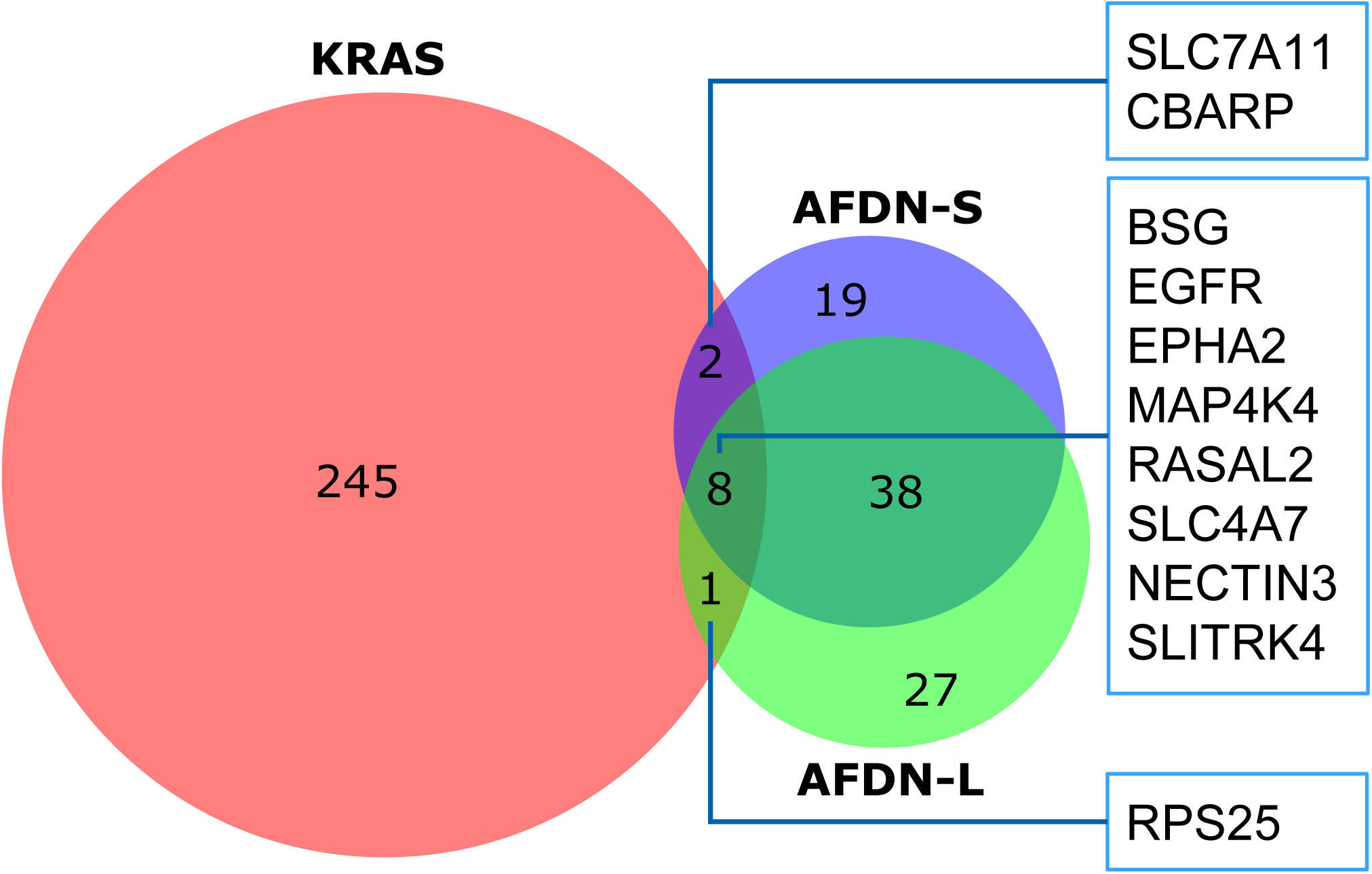
Venn diagram of the numbers of shared and unique proteins between the dynamic KRAS interactome identified by our BioID study and the published interactome of AFDN isoforms

**Supplementary Fig 6.**
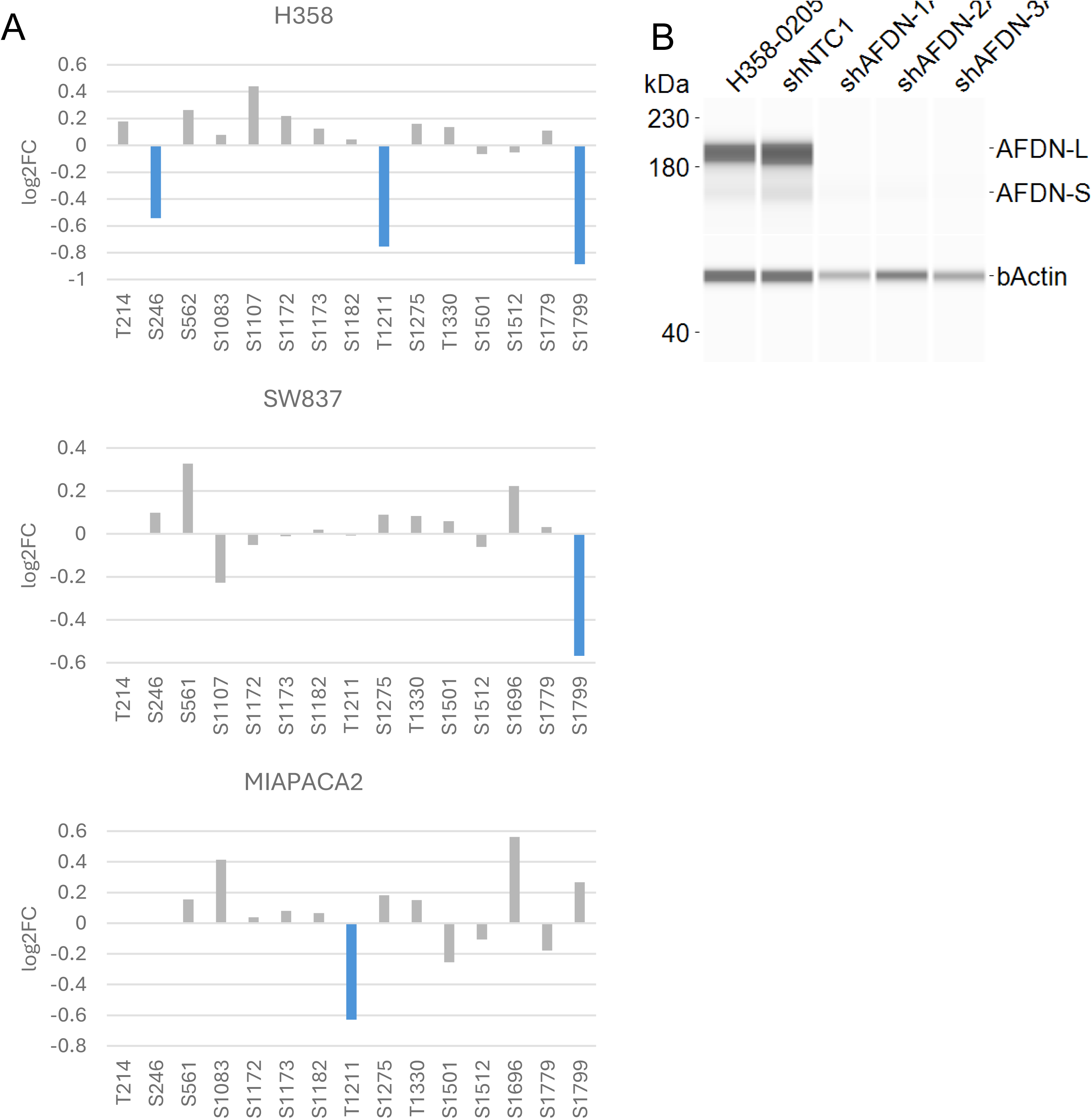
(A) Phosphorylation of different serine residue of AFDN in three cancer cell lines treated with KRASG12C inhibitor MRTX1257 based on phosphoproteome data retrieved from Jennifer Klomp et al.’s study. Significant fold-change (adjusted p-value < 0.05) were highlighted in blue **(B)** shRNA-induced AFDN knockdown in H358 cells. NTC: non-targeting control

**Supplementary Fig 7.**
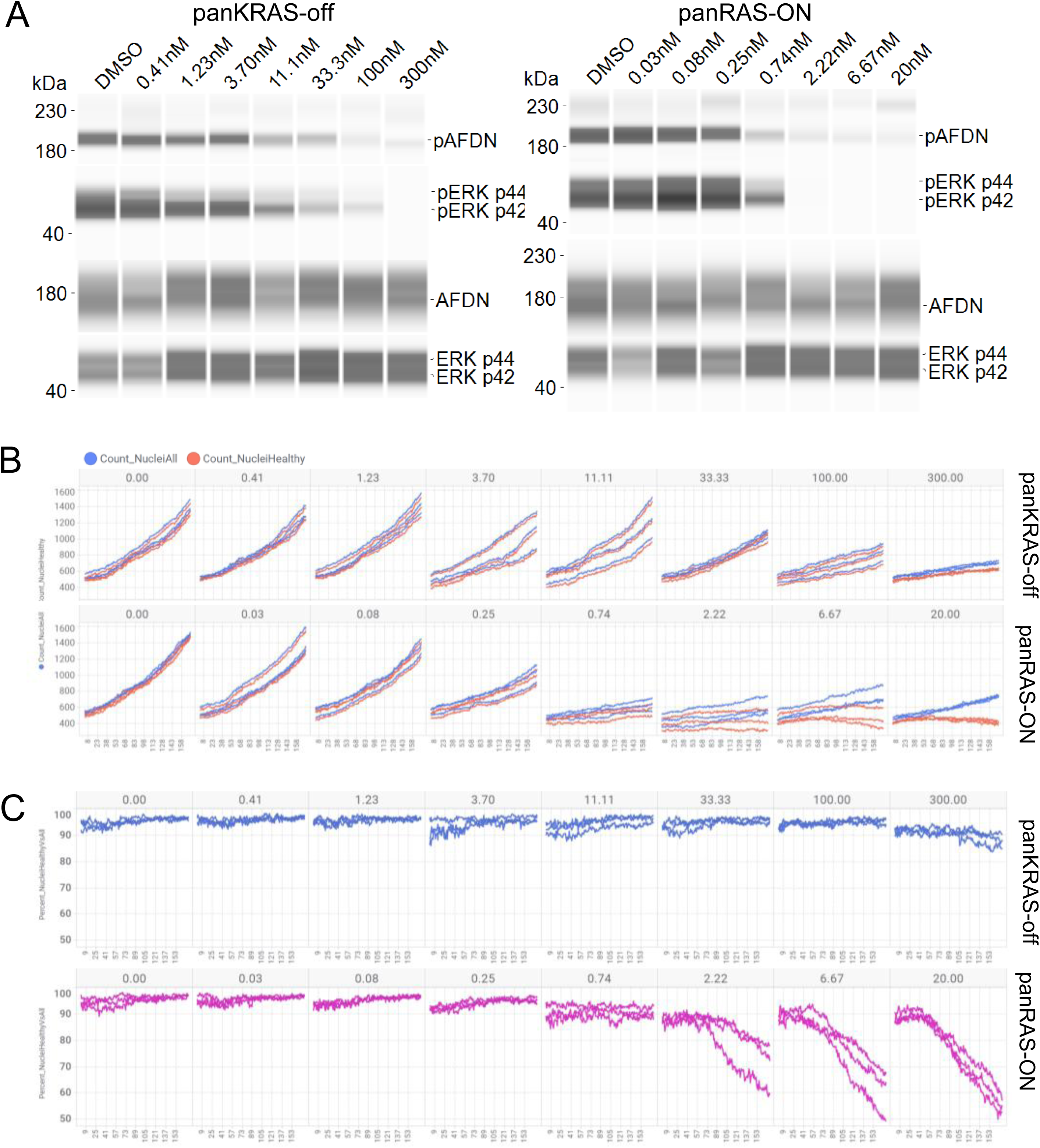
(A) Immunoblot of phosphorylated p-ERK and p-AFDN in H358 cells treated with different concentrations of panRAS-ON and panKRAS-off. **(B)** The number of healthy nuclei and total nuclei of H358 cells treated with different concentrations of panRAS-ON (bottom) and panKRAS-off (top) over 48 hours. Blue: total nuclei. Red: healthy nuclei. **(C)** Percentage of healthy nuclei from H358 cells treated with different concentrations of panRAS-ON (bottom) or panKRAS-off (top) over 48 hours.

**Supplementary Fig 8.**
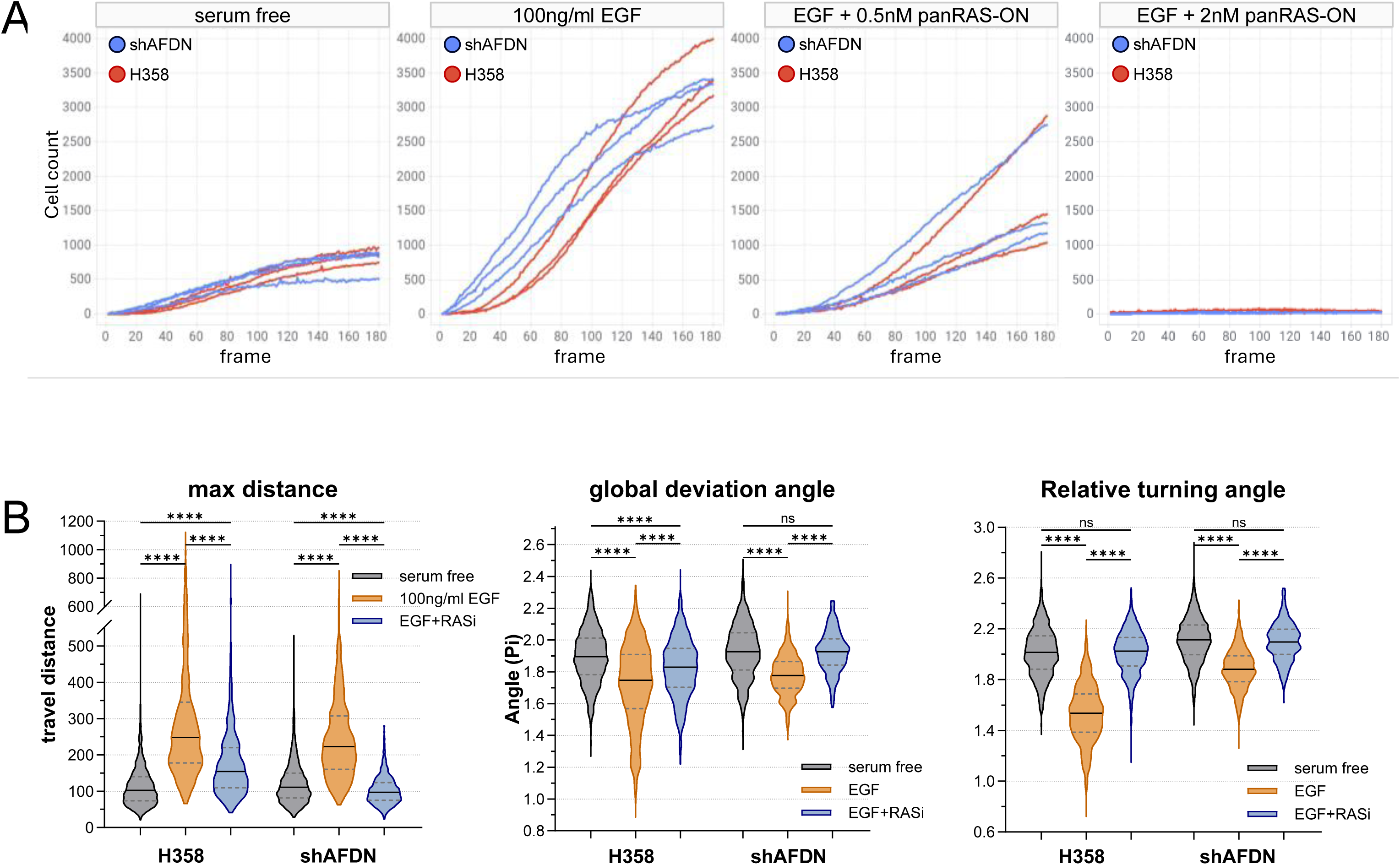
(A) Number of cells entering the wound area over the entire 48 hours of wound healing process between H358 cells and the shAFDN-knockdown cells under different conditions. Both cell lines were grown in serum-free media during the wound healing process supplemented with 100 ng/ml EGF or additional panRAS-ON (0.5 nM or 2 nM) in triplicate. **(B)** Track measures calculated from aggregated individual cell tracking. Parameters were calculated using equations from Svensson et al (15). Max distance: the farthest distance away from the beginning location of individual cell during its entire migration track. Global deviation angle: the degree of a track deviates from the upright direction. Relative turning angle: the degree of a track deviates from its previous direction. (✱: FDR<0.05; ✱✱: FDR<0.005; ✱✱✱: FDR<0.0005; ✱✱✱✱: FDR<0.0001)

**Supplementary Fig 9.**
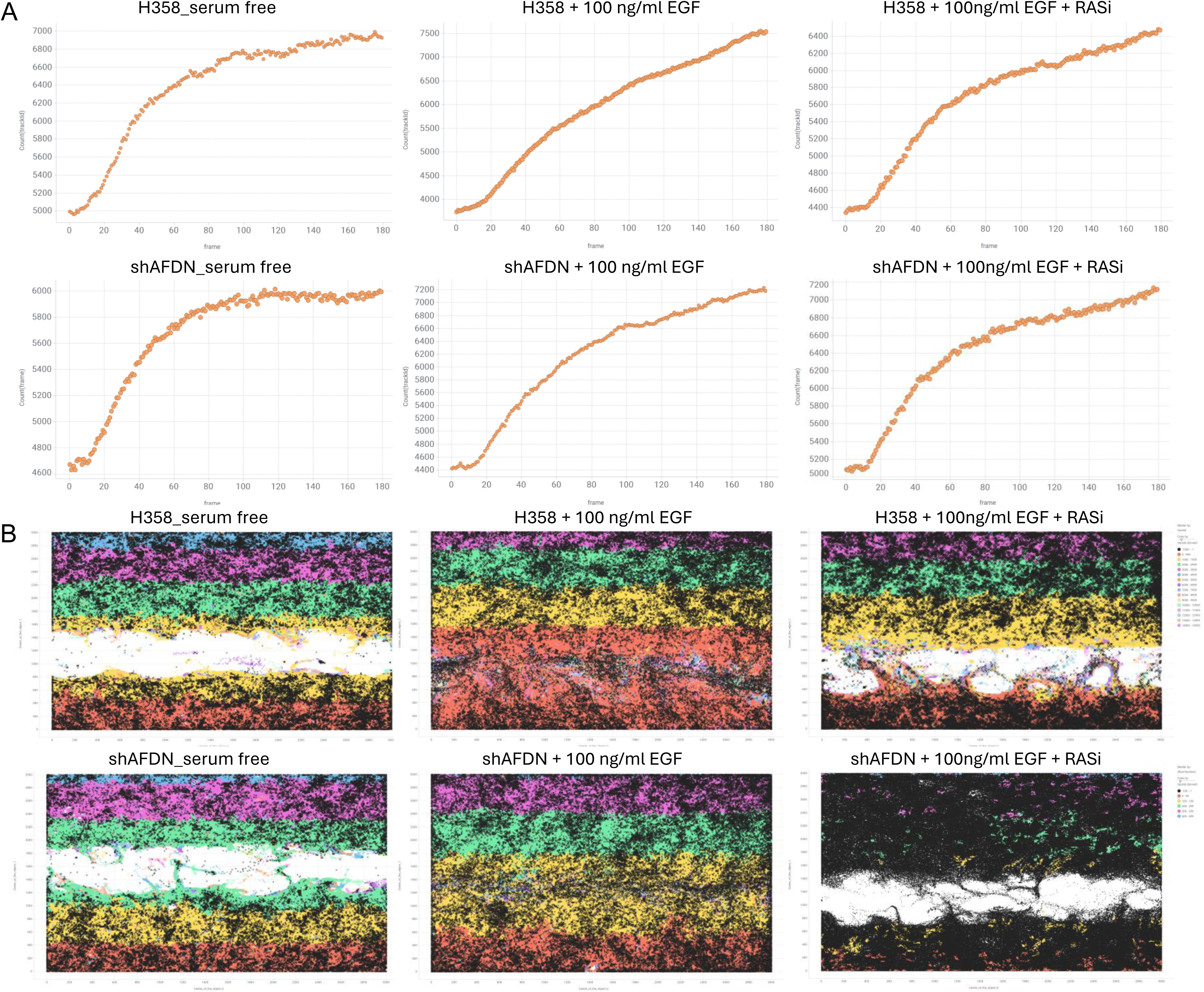
(A) Number of total segmented individual cells of both H358 and shAFDN-knockdown cell lines identified from imaging per time point over the 48 hours of wound healing process. (B) Location of all segmented individual cells identified from imaging per time point over the 48 hours of wound healing process. Trackable cells between frames were colored differently based on initial location. Untraceable cells were colored in black.

**Supplementary Fig 10.**
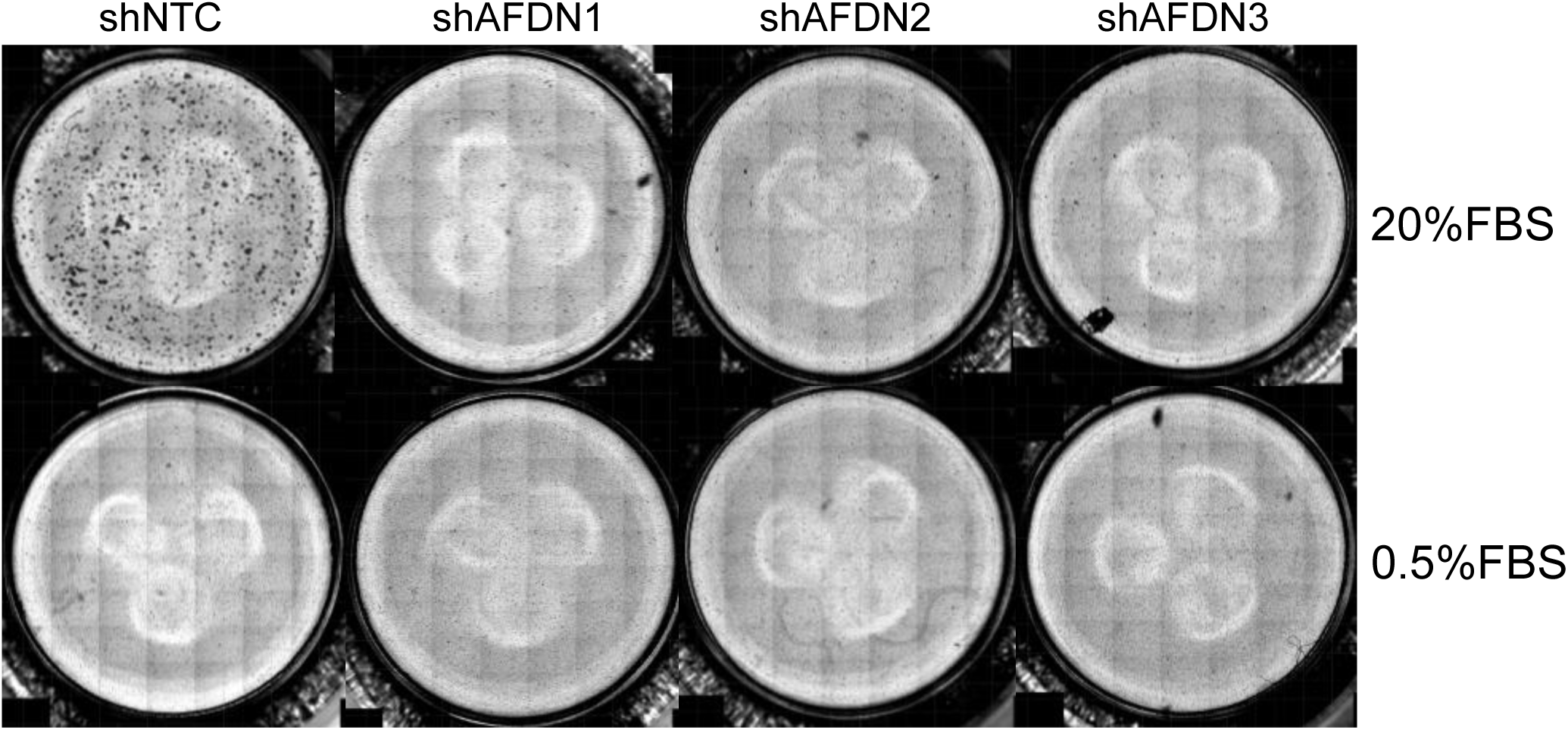
Transwell migration assay of H358 cells treated with shAFDN or non-targeting control. 20% FBS was used as chemoattractant

