## Supplementary File 1 for "Remodeling of KRAS interactome induced by clinically relevant RAS inhibitors reveals convergent responses and a KRAS-mediated regulation of directional cell migration"

#### TurboID-KRAS4A^G12D^

CTACGTTTAAACGCCACCATGGACTACAAAGACCATGACGGTGATTATAAAGATCATGACATCGATTACAAGGATGACGATGACAAGGGAGGCTCTGGTAAAGACAATACTGTGCCTCTGAAGCTGATCGCTCTCCTGGCTAATGGCGAGTTCCATAGTGGCGAACAGCTGGGAGAAACCCTGGGCATGTCCAGGGCCGCTATCAACAAGCACATTCAGACTCTGCGCGACTGGGGCGTGGACGTGTTCACCGTGCCCGGAAAGGGCTACTCTCTGCCCGAGCCTATCCCGCTGCTGAACGCTAAACAGATTCTGGGACAGCTGGACGGCGGGAGCGTGGCAGTCCTGCCTGTGGTCGACTCCACCAATCAGTACCTGCTGGATCGAATCGGCGAGCTGAAGAGTGGGGATGCTTGCATTGCAGAATATCAGCAGGCAGGGAGAGGAAGCAGAGGGAGGAAATGGTTCTCTCCTTTTGGAGCTAACCTGTACCTGAGTATGTTTTGGCGCCTGAAGCGGGGACCAGCAGCAATCGGCCTGGGCCCGGTCATCGGAATTGTCATGGCAGAAGCGCTGCGAAAGCTGGGAGCAGACAAGGTGCGAGTCAAATGGCCCAATGACCTGTATCTGCAGGATAGAAAGCTGGCAGGCATCCTGGTGGAGCTGGCCGGAATAACAGGCGATGCTGCACAGATCGTCATTGGCGCCGGGATTAACGTGGCTATGAGGCGCGTGGAGGAAAGCGTGGTCAATCAGGGCTGGATCACACTGCAGGAAGCAGGGATTAACCTGGACAGGAATACTCTGGCCGCTACGCTGATCCGAGAGCTGCGGGCAGCCCTGGAACTGTTCGAGCAGGAAGGCCTGGCTCCATATCTGCCACGGTGGGAGAAGCTGGATAACTTCATCAATAGACCCGTGAAGCTGATCATTGGGGACAAAGAGATTTTCGGGATTAGCCGGGGGATTGATAAACAGGGAGCCCTGCTGCTGGAACAGGACGGAGTTATCAAACCCTGGATGGGCGGAGAAATCAGTCTGCGGTCTGCCGAAAAGGGATCCAGCGGCGGCGGCAGCGGCGGCGGCGGCAGCACTGAATATAAACTTGTGGTAGTTGGAGCTGACGGCGTAGGCAAGAGTGCCTTGACGATACAGCTAATTCAGAATCATTTTGTGGACGAATATGATCCAACAATAGAGGATTCCTACAGGAAGCAAGTAGTAATTGATGGAGAAACCTGTCTCTTGGATATTCTCGACACAGCAGGTCAAGAGGAGTACAGTGCAATGAGGGACCAGTACATGAGGACTGGGGAGGGCTTTCTTTGTGTATTTGCCATAAATAATACTAAATCATTTGAAGATATTCACCATTATAGAGAACAAATTAAAAGAGTTAAGGACTCTGAAGATGTACCTATGGTCCTAGTAGGAAATAAATGTGATTTGCCTTCTAGAACAGTAGACACAAAACAGGCTCAGGACTTAGCAAGAAGTTATGGAATTCCTTTTATTGAAACATCAGCAAAGACAAGACAGAGAGTGGAGGATGCTTTTTATACATTGGTGAGAGAGATCCGACAATACAGATTGAAAAAAATCAGCAAAGAAGAAAAGACTCCTGGCTGTGTGAAAATTAAAAAATGCATTATAATGTAGTAAGCTAGCGGAGTTCCGCG

#### TurboID-KRAS4A^G12V^

CTACGTTTAAACGCCACCATGGACTACAAAGACCATGACGGTGATTATAAAGATCATGACATCGATTACAAGGATGACGATGACAAGGGAGGCTCTGGTAAAGACAATACTGTGCCTCTGAAGCTGATCGCTCTCCTGGCTAATGGCGAGTTCCATAGTGGCGAACAGCTGGGAGAAACCCTGGGCATGTCCAGGGCCGCTATCAACAAGCACATTCAGACTCTGCGCGACTGGGGCGTGGACGTGTTCACCGTGCCCGGAAAGGGCTACTCTCTGCCCGAGCCTATCCCGCTGCTGAACGCTAAACAGATTCTGGGACAGCTGGACGGCGGGAGCGTGGCAGTCCTGCCTGTGGTCGACTCCACCAATCAGTACCTGCTGGATCGAATCGGCGAGCTGAAGAGTGGGGATGCTTGCATTGCAGAATATCAGCAGGCAGGGAGAGGAAGCAGAGGGAGGAAATGGTTCTCTCCTTTTGGAGCTAACCTGTACCTGAGTATGTTTTGGCGCCTGAAGCGGGGACCAGCAGCAATCGGCCTGGGCCCGGTCATCGGAATTGTCATGGCAGAAGCGCTGCGAAAGCTGGGAGCAGACAAGGTGCGAGTCAAATGGCCCAATGACCTGTATCTGCAGGATAGAAAGCTGGCAGGCATCCTGGTGGAGCTGGCCGGAATAACAGGCGATGCTGCACAGATCGTCATTGGCGCCGGGATTAACGTGGCTATGAGGCGCGTGGAGGAAAGCGTGGTCAATCAGGGCTGGATCACACTGCAGGAAGCAGGGATTAACCTGGACAGGAATACTCTGGCCGCTACGCTGATCCGAGAGCTGCGGGCAGCCCTGGAACTGTTCGAGCAGGAAGGCCTGGCTCCATATCTGCCACGGTGGGAGAAGCTGGATAACTTCATCAATAGACCCGTGAAGCTGATCATTGGGGACAAAGAGATTTTCGGGATTAGCCGGGGGATTGATAAACAGGGAGCCCTGCTGCTGGAACAGGACGGAGTTATCAAACCCTGGATGGGCGGAGAAATCAGTCTGCGGTCTGCCGAAAAGGGATCCAGCGGCGGCGGCAGCGGCGGCGGCGGCAGCACTGAATATAAACTTGTGGTAGTTGGAGCTGTTGGCGTAGGCAAGAGTGCCTTGACGATACAGCTAATTCAGAATCATTTTGTGGACGAATATGATCCAACAATAGAGGATTCCTACAGGAAGCAAGTAGTAATTGATGGAGAAACCTGTCTCTTGGATATTCTCGACACAGCAGGTCAAGAGGAGTACAGTGCAATGAGGGACCAGTACATGAGGACTGGGGAGGGCTTTCTTTGTGTATTTGCCATAAATAATACTAAATCATTTGAAGATATTCACCATTATAGAGAACAAATTAAAAGAGTTAAGGACTCTGAAGATGTACCTATGGTCCTAGTAGGAAATAAATGTGATTTGCCTTCTAGAACAGTAGACACAAAACAGGCTCAGGACTTAGCAAGAAGTTATGGAATTCCTTTTATTGAAACATCAGCAAAGACAAGACAGAGAGTGGAGGATGCTTTTTATACATTGGTGAGAGAGATCCGACAATACAGATTGAAAAAAATCAGCAAAGAAGAAAAGACTCCTGGCTGTGTGAAAATTAAAAAATGCATTATAATGTAGTAAGCTAGCGGAGTTCCGCG

#### TurboID-KRAS4A^WT^

CTACGTTTAAACGCCACCATGGACTACAAAGACCATGACGGTGATTATAAAGATCATGACATCGATTACAAGGATGACGATGACAAGGGAGGCTCTGGTAAAGACAATACTGTGCCTCTGAAGCTGATCGCTCTCCTGGCTAATGGCGAGTTCCATAGTGGCGAACAGCTGGGAGAAACCCTGGGCATGTCCAGGGCCGCTATCAACAAGCACATTCAGACTCTGCGCGACTGGGGCGTGGACGTGTTCACCGTGCCCGGAAAGGGCTACTCTCTGCCCGAGCCTATCCCGCTGCTGAACGCTAAACAGATTCTGGGACAGCTGGACGGCGGGAGCGTGGCAGTCCTGCCTGTGGTCGACTCCACCAATCAGTACCTGCTGGATCGAATCGGCGAGCTGAAGAGTGGGGATGCTTGCATTGCAGAATATCAGCAGGCAGGGAGAGGAAGCAGAGGGAGGAAATGGTTCTCTCCTTTTGGAGCTAACCTGTACCTGAGTATGTTTTGGCGCCTGAAGCGGGGACCAGCAGCAATCGGCCTGGGCCCGGTCATCGGAATTGTCATGGCAGAAGCGCTGCGAAAGCTGGGAGCAGACAAGGTGCGAGTCAAATGGCCCAATGACCTGTATCTGCAGGATAGAAAGCTGGCAGGCATCCTGGTGGAGCTGGCCGGAATAACAGGCGATGCTGCACAGATCGTCATTGGCGCCGGGATTAACGTGGCTATGAGGCGCGTGGAGGAAAGCGTGGTCAATCAGGGCTGGATCACACTGCAGGAAGCAGGGATTAACCTGGACAGGAATACTCTGGCCGCTACGCTGATCCGAGAGCTGCGGGCAGCCCTGGAACTGTTCGAGCAGGAAGGCCTGGCTCCATATCTGCCACGGTGGGAGAAGCTGGATAACTTCATCAATAGACCCGTGAAGCTGATCATTGGGGACAAAGAGATTTTCGGGATTAGCCGGGGGATTGATAAACAGGGAGCCCTGCTGCTGGAACAGGACGGAGTTATCAAACCCTGGATGGGCGGAGAAATCAGTCTGCGGTCTGCCGAAAAGGGATCCAGCGGCGGCGGCAGCGGCGGCGGCGGCAGCACTGAATATAAACTTGTGGTAGTTGGAGCTGGTGGCGTAGGCAAGAGTGCCTTGACGATACAGCTAATTCAGAATCATTTTGTGGACGAATATGATCCAACAATAGAGGATTCCTACAGGAAGCAAGTAGTAATTGATGGAGAAACCTGTCTCTTGGATATTCTCGACACAGCAGGTCAAGAGGAGTACAGTGCAATGAGGGACCAGTACATGAGGACTGGGGAGGGCTTTCTTTGTGTATTTGCCATAAATAATACTAAATCATTTGAAGATATTCACCATTATAGAGAACAAATTAAAAGAGTTAAGGACTCTGAAGATGTACCTATGGTCCTAGTAGGAAATAAATGTGATTTGCCTTCTAGAACAGTAGACACAAAACAGGCTCAGGACTTAGCAAGAAGTTATGGAATTCCTTTTATTGAAACATCAGCAAAGACAAGACAGAGAGTGGAGGATGCTTTTTATACATTGGTGAGAGAGATCCGACAATACAGATTGAAAAAAATCAGCAAAGAAGAAAAGACTCCTGGCTGTGTGAAAATTAAAAAATGCATTATAATGTAGTAAGCTAGCGGAGTTCCGCG

#### TurboID-KRAS4B^G12D^

CTACGTTTAAACGCCACCATGGACTACAAAGACCATGACGGTGATTATAAAGATCATGACATCGATTACAAGGATGACGATGACAAGGGAGGCTCTGGTAAAGACAATACTGTGCCTCTGAAGCTGATCGCTCTCCTGGCTAATGGCGAGTTCCATAGTGGCGAACAGCTGGGAGAAACCCTGGGCATGTCCAGGGCCGCTATCAACAAGCACATTCAGACTCTGCGCGACTGGGGCGTGGACGTGTTCACCGTGCCCGGAAAGGGCTACTCTCTGCCCGAGCCTATCCCGCTGCTGAACGCTAAACAGATTCTGGGACAGCTGGACGGCGGGAGCGTGGCAGTCCTGCCTGTGGTCGACTCCACCAATCAGTACCTGCTGGATCGAATCGGCGAGCTGAAGAGTGGGGATGCTTGCATTGCAGAATATCAGCAGGCAGGGAGAGGAAGCAGAGGGAGGAAATGGTTCTCTCCTTTTGGAGCTAACCTGTACCTGAGTATGTTTTGGCGCCTGAAGCGGGGACCAGCAGCAATCGGCCTGGGCCCGGTCATCGGAATTGTCATGGCAGAAGCGCTGCGAAAGCTGGGAGCAGACAAGGTGCGAGTCAAATGGCCCAATGACCTGTATCTGCAGGATAGAAAGCTGGCAGGCATCCTGGTGGAGCTGGCCGGAATAACAGGCGATGCTGCACAGATCGTCATTGGCGCCGGGATTAACGTGGCTATGAGGCGCGTGGAGGAAAGCGTGGTCAATCAGGGCTGGATCACACTGCAGGAAGCAGGGATTAACCTGGACAGGAATACTCTGGCCGCTACGCTGATCCGAGAGCTGCGGGCAGCCCTGGAACTGTTCGAGCAGGAAGGCCTGGCTCCATATCTGCCACGGTGGGAGAAGCTGGATAACTTCATCAATAGACCCGTGAAGCTGATCATTGGGGACAAAGAGATTTTCGGGATTAGCCGGGGGATTGATAAACAGGGAGCCCTGCTGCTGGAACAGGACGGAGTTATCAAACCCTGGATGGGCGGAGAAATCAGTCTGCGGTCTGCCGAAAAGGGATCCAGCGGCGGCGGCAGCGGCGGCGGCGGCAGCACTGAATATAAACTTGTGGTAGTTGGAGCTGACGGCGTAGGCAAGAGTGCCTTGACGATACAGCTAATTCAGAATCATTTTGTGGACGAATATGATCCAACAATAGAGGATTCCTACAGGAAGCAAGTAGTAATTGATGGAGAAACCTGTCTCTTGGATATTCTCGACACAGCAGGTCAAGAGGAGTACAGTGCAATGAGGGACCAGTACATGAGGACTGGGGAGGGCTTTCTTTGTGTATTTGCCATAAATAATACTAAATCATTTGAAGATATTCACCATTATAGAGAACAAATTAAAAGAGTTAAGGACTCTGAAGATGTACCTATGGTCCTAGTAGGAAATAAATGTGATTTGCCTTCTAGAACAGTAGACACAAAACAGGCTCAGGACTTAGCAAGAAGTTATGGAATTCCTTTTATTGAAACATCAGCAAAGACAAGACAGGGTGTTGATGATGCCTTCTATACATTAGTTCGAGAAATTCGAAAACATAAAGAAAAGATGAGCAAAGATGGTAAAAAGAAGAAAAAGAAGTCAAAGACAAAGTGTGTAATTATGTAGTAAGCTAGCGGAGTTCCGCG

#### TurboID-KRAS4B^G12V^

CTACGTTTAAACGCCACCATGGACTACAAAGACCATGACGGTGATTATAAAGATCATGACATCGATTACAAGGATGACGATGACAAGGGAGGCTCTGGTAAAGACAATACTGTGCCTCTGAAGCTGATCGCTCTCCTGGCTAATGGCGAGTTCCATAGTGGCGAACAGCTGGGAGAAACCCTGGGCATGTCCAGGGCCGCTATCAACAAGCACATTCAGACTCTGCGCGACTGGGGCGTGGACGTGTTCACCGTGCCCGGAAAGGGCTACTCTCTGCCCGAGCCTATCCCGCTGCTGAACGCTAAACAGATTCTGGGACAGCTGGACGGCGGGAGCGTGGCAGTCCTGCCTGTGGTCGACTCCACCAATCAGTACCTGCTGGATCGAATCGGCGAGCTGAAGAGTGGGGATGCTTGCATTGCAGAATATCAGCAGGCAGGGAGAGGAAGCAGAGGGAGGAAATGGTTCTCTCCTTTTGGAGCTAACCTGTACCTGAGTATGTTTTGGCGCCTGAAGCGGGGACCAGCAGCAATCGGCCTGGGCCCGGTCATCGGAATTGTCATGGCAGAAGCGCTGCGAAAGCTGGGAGCAGACAAGGTGCGAGTCAAATGGCCCAATGACCTGTATCTGCAGGATAGAAAGCTGGCAGGCATCCTGGTGGAGCTGGCCGGAATAACAGGCGATGCTGCACAGATCGTCATTGGCGCCGGGATTAACGTGGCTATGAGGCGCGTGGAGGAAAGCGTGGTCAATCAGGGCTGGATCACACTGCAGGAAGCAGGGATTAACCTGGACAGGAATACTCTGGCCGCTACGCTGATCCGAGAGCTGCGGGCAGCCCTGGAACTGTTCGAGCAGGAAGGCCTGGCTCCATATCTGCCACGGTGGGAGAAGCTGGATAACTTCATCAATAGACCCGTGAAGCTGATCATTGGGGACAAAGAGATTTTCGGGATTAGCCGGGGGATTGATAAACAGGGAGCCCTGCTGCTGGAACAGGACGGAGTTATCAAACCCTGGATGGGCGGAGAAATCAGTCTGCGGTCTGCCGAAAAGGGATCCAGCGGCGGCGGCAGCGGCGGCGGCGGCAGCACTGAATATAAACTTGTGGTAGTTGGAGCTGTTGGCGTAGGCAAGAGTGCCTTGACGATACAGCTAATTCAGAATCATTTTGTGGACGAATATGATCCAACAATAGAGGATTCCTACAGGAAGCAAGTAGTAATTGATGGAGAAACCTGTCTCTTGGATATTCTCGACACAGCAGGTCAAGAGGAGTACAGTGCAATGAGGGACCAGTACATGAGGACTGGGGAGGGCTTTCTTTGTGTATTTGCCATAAATAATACTAAATCATTTGAAGATATTCACCATTATAGAGAACAAATTAAAAGAGTTAAGGACTCTGAAGATGTACCTATGGTCCTAGTAGGAAATAAATGTGATTTGCCTTCTAGAACAGTAGACACAAAACAGGCTCAGGACTTAGCAAGAAGTTATGGAATTCCTTTTATTGAAACATCAGCAAAGACAAGACAGGGTGTTGATGATGCCTTCTATACATTAGTTCGAGAAATTCGAAAACATAAAGAAAAGATGAGCAAAGATGGTAAAAAGAAGAAAAAGAAGTCAAAGACAAAGTGTGTAATTATGTAGTAAGCTAGCGGAGTTCCGCG

#### TurboID-KRAS4B^WT^

CTACGTTTAAACGCCACCATGGACTACAAAGACCATGACGGTGATTATAAAGATCATGACATCGATTACAAGGATGACGATGACAAGGGAGGCTCTGGTAAAGACAATACTGTGCCTCTGAAGCTGATCGCTCTCCTGGCTAATGGCGAGTTCCATAGTGGCGAACAGCTGGGAGAAACCCTGGGCATGTCCAGGGCCGCTATCAACAAGCACATTCAGACTCTGCGCGACTGGGGCGTGGACGTGTTCACCGTGCCCGGAAAGGGCTACTCTCTGCCCGAGCCTATCCCGCTGCTGAACGCTAAACAGATTCTGGGACAGCTGGACGGCGGGAGCGTGGCAGTCCTGCCTGTGGTCGACTCCACCAATCAGTACCTGCTGGATCGAATCGGCGAGCTGAAGAGTGGGGATGCTTGCATTGCAGAATATCAGCAGGCAGGGAGAGGAAGCAGAGGGAGGAAATGGTTCTCTCCTTTTGGAGCTAACCTGTACCTGAGTATGTTTTGGCGCCTGAAGCGGGGACCAGCAGCAATCGGCCTGGGCCCGGTCATCGGAATTGTCATGGCAGAAGCGCTGCGAAAGCTGGGAGCAGACAAGGTGCGAGTCAAATGGCCCAATGACCTGTATCTGCAGGATAGAAAGCTGGCAGGCATCCTGGTGGAGCTGGCCGGAATAACAGGCGATGCTGCACAGATCGTCATTGGCGCCGGGATTAACGTGGCTATGAGGCGCGTGGAGGAAAGCGTGGTCAATCAGGGCTGGATCACACTGCAGGAAGCAGGGATTAACCTGGACAGGAATACTCTGGCCGCTACGCTGATCCGAGAGCTGCGGGCAGCCCTGGAACTGTTCGAGCAGGAAGGCCTGGCTCCATATCTGCCACGGTGGGAGAAGCTGGATAACTTCATCAATAGACCCGTGAAGCTGATCATTGGGGACAAAGAGATTTTCGGGATTAGCCGGGGGATTGATAAACAGGGAGCCCTGCTGCTGGAACAGGACGGAGTTATCAAACCCTGGATGGGCGGAGAAATCAGTCTGCGGTCTGCCGAAAAGGGATCCAGCGGCGGCGGCAGCGGCGGCGGCGGCAGCACTGAATATAAACTTGTGGTAGTTGGAGCTGGTGGCGTAGGCAAGAGTGCCTTGACGATACAGCTAATTCAGAATCATTTTGTGGACGAATATGATCCAACAATAGAGGATTCCTACAGGAAGCAAGTAGTAATTGATGGAGAAACCTGTCTCTTGGATATTCTCGACACAGCAGGTCAAGAGGAGTACAGTGCAATGAGGGACCAGTACATGAGGACTGGGGAGGGCTTTCTTTGTGTATTTGCCATAAATAATACTAAATCATTTGAAGATATTCACCATTATAGAGAACAAATTAAAAGAGTTAAGGACTCTGAAGATGTACCTATGGTCCTAGTAGGAAATAAATGTGATTTGCCTTCTAGAACAGTAGACACAAAACAGGCTCAGGACTTAGCAAGAAGTTATGGAATTCCTTTTATTGAAACATCAGCAAAGACAAGACAGGGTGTTGATGATGCCTTCTATACATTAGTTCGAGAAATTCGAAAACATAAAGAAAAGATGAGCAAAGATGGTAAAAAGAAGAAAAAGAAGTCAAAGACAAAGTGTGTAATTATGTAGTAAGCTAGCGGAGTTCCGCG

#### RA^KO^-AFDN

CATCATCACCATCATCACCATATACCCAAAAAAACAAAGAAGCATCTGGAGGGCAAGACTCCTAAGGGAAAAGAAAGAGCTGATGGAAGTGGCTACGGCTCCACTCTGCCTCCTGAGAAGCTGCCCTACCTGGTCGAGCTGTCCCCTGGACGCAGAAATCATTTCGCATACTACAATTATCACACCTACGAAGATGGAAGCGATAGCAGAGATAAGCCCAAGCTGTACCGGTTGCAGCTCAGTGTGACCGAAGTCGGAACAGAAAAGCTCGACGATAACTCTATCCAGCTGTTTGGGCCCGGAATCCAGCCGCACCACTGTGATCTGACCAATATGGACGGCGTAGTGACTGTGACCCCACGCAGCATGGATGCAGAAACTTATGTAGAGGGACAGCGCATTTCCGAAACGACAATGCTGCAATCCGGAATGAAAGTCCAGTTTGGAGCTAGCCATGTCTTCAAGTTCGTGGACCCCAGTCAGGACCACGCTCTGGCCAAACGCTCTGTGGACGGAGGACTCATGGTGAAGGGACCCAGGCACAAGCCAGGTATCGTCCAGGAGACAACTTTCGATTTGGGCGGTGACATCCACAGCGGCACTGCGCTGCCAACTTCCAAATCAACCACCCGGCTTGACTCAGATAGGGTGTCCTCAGCGAGTTCAACCGCTGAGCGGGGGATGGTCAAGCCAATGATAAGAGTGGAGCAGCAGCCAGACTATCGGCGGCAGGAGAGCCGAACACAAGACGCCAGCGGCCCTGAACTGATACTGCCCGCCAGCATTGAGTTTCGGGAATCTTCTGAAGATTCTTTTCTTTCCGCCATTATTAACTACACTAACTCCTCTACCGTCCATTTCAAGCTGTCCCCAACCTACGTGCTTTACATGGCCTGCCGGTACGTGTTGAGCAACCAATACAGGCCGGATATATCCCCAACCGAGAGGACGCACAAAGTGATCGCTGTGGTCAACAAGATGGTCTCCATGATGGAGGGCGTGATACAGAAGCAAAAGAACATTGCCGGAGCCCTCGCCTTTTGGATGGCGAACGCAAGCGAGCTGCTGAACTTCATCAAACAGGATAGAGACCTTTCTAGGATCACCCTGGATGCCCAGGATGTCCTCGCTCACCTGGTGCAAATGGCCTTTAAGTATTTGGTGCATTGTCTTCAAAGTGAACTTAACAACTATATGCCAGCTTTCCTTGACGACCCAGAAGAAAATTCACTGCAACGCCCTAAGATTGATGACGTGTTGCACACCCTCACCGGCGCGATGAGCCTGCTGCGGAGATGCAGGGTGAACGCAGCCTTGACCATACAGCTGTTTAGCCAGCTGTTCCATTTCATTAATATGTGGCTGTTCAACAGACTCGTGACTGACCCAGACAGCGGGCTGTGCTCTCACTACTGGGGGGCTATCATCAGGCAGCAACTCGGGCACATTGAGGCCTGGGCCGAAAAGCAGGGCCTGGAACTTGCTGCCGACTGTCACCTGTCTAGAATCGTCCAGGCCACAACACTGCTGACCATGGATAAGTATGCCCCTGATGACATTCCCAACATCAATTCTACGTGTTTCAAGTTGAACTCACTCCAACTCCAAGCCCTTCTGCAGAACTACCATTGCGCCCCCGATGAGCCATTCATACCCACAGACCTGATTGAAAACGTCGTCACTGTGGCTGAGAACACAGCAGACGAGCTGGCACGATCAGATGGCCGGGAAGTGCAGCTTGAGGAGGACCCAGATCTGCAGCTGCCTTTTCTTCTTCCAGAAGACGGTTATAGCTGTGATGTCGTCCGCAATATCCCCAATGGGCTCCAAGAATTTTTGGACCCCCTGTGCCAGAGAGGGTTCTGCCGACTGATCCCTCACACGCGAAGCCCGGGGACCTGGACCATATACTTCGAGGGCGCTGACTACGAATCCCACCTGTTGCGGGAAAATACTGAACTGGCCCAACCACTGCGGAAGGAACCCGAGATCATTACAGTTACGTTGAAAAAGCAGAACGGAATGGGCCTGTCTATAGTTGCCGCCAAAGGCGCGGGTCAAGACAAACTTGGAATTTATGTCAAGAGCGTCGTTAAAGGCGGTGCAGCAGATGTGGACGGAAGATTGGCAGCCGGCGACCAGCTGCTTTCTGTAGACGGCCGCAGTCTGGTGGGGCTGTCCCAGGAACGAGCCGCGGAGCTGATGACCAGAACTTCAAGTGTGGTTACCCTTGAGGTAGCCAAGCAGGGGGCTATTTATCACGGGCTTGCCACCCTGCTCAATCAGCCATCTCCTATGATGCAGAGGATATCCGATCGGAGGGGCTCTGGCAAGCCAAGACCTAAGTCTGAGGGATTCGAATTGTATAACAATTCAACCCAAAACGGTTCACCAGAGAGCCCACAGCTGCCCTGGGCAGAGTATAGCGAACCAAAAAAACTGCCCGGGGATGACCGGTTGATGAAAAACCGAGCAGACCACCGATCATCTCCTAATGTTGCTAATCAGCCCCCCTCACCAGGGGGCAAATCTGCATACGCCAGTGGAACTACAGCAAAAATTACCAGTGTCAGTACCGGGAATCTGTGTACTGAGGAGCAGACTCCCCCCCCCCGACCCGAAGCCTATCCAATCCCCACTCAGACTTACACACGGGAGTACTTTACTTTCCCGGCCAGTAAGTCACAAGACAGAATGGCCCCTCCTCAAAACCAATGGCCAAACTATGAAGAGAAGCCGCATATGCACACGGATTCCAACCACAGCTCTATTGCCATACAGAGGGTTACCAGGAGTCAAGAAGAGCTGAGAGAAGACAAGGCATACCAGCTGGAGAGGCACCGGATAGAAGCAGCTATGGATCGGAAAAGCGACAGCGACATGTGGATCAACCAAAGCTCATCACTGGACTCCTCCACCTCCAGCCAAGAACATCTCAATCACAGCAGCAAGTCCGTCACGCCAGCCAGTACCCTGACTAAGAGTGGACCAGGACGCTGGAAGACCCCCGCCGCTATCCCCGCTACACCCGTGGCGGTGTCTCAACCTATCAGGACCGACCTGCCACCACCCCCCCCCCCTCCCCCAGTGCATTACGCCGGGGACTTCGATGGGATGAGCATGGACCTCCCCTTGCCACCACCTCCTAGTGCCAATCAGATCGGTCTCCCATCCGCGCAGGTGGCCGCTGCTGAACGGCGAAAGCGCGAAGAACACCAGCGATGGTATGAGAAGGAGAAGGCCCGCCTGGAGGAAGAGCGAGAAAGAAAGCGCCGCGAGCAAGAGCGGAAACTTGGTCAAATGCGAACTCAATCTCTGAATCCGGCTCCATTCTCACCGCTGACAGCTCAGCAGATGAAGCCCGAAAAGCCCTCCACCCTCCAGCGCCCCCAAGAGACCGTGATCCGGGAGCTCCAGCCTCAGCAGCAGCCACGAACAATCGAACGCAGGGACCTTCAGTACATTACAGTCTCTAAAGAAGAACTGAGTTCAGGGGATAGTCTGTCTCCCGACCCATGGAAGCGGGATGCAAAGGAGAAGTTGGAGAAGCAGCAGCAGATGCACATTGTCGATATGCTCAGCAAGGAGATACAGGAGCTTCAGAGCAAGCCGGATAGATCCGCCGAAGAAAGCGACCGCCTCCGGAAACTGATGCTCGAATGGCAGTTTCAGAAGCGATTGCAAGAGAGCAAGCAGAAGGACGAAGACGACGAGGAGGAAGAGGACGATGATGTGGATACCATGCTTATTATGCAGCGCCTGGAAGCCGAAAGACGCGCACGGCTGCAGGACGAGGAACGCCGCAGGCAGCAGCAGCTGGAAGAGATGAGGAAACGAGAAGCTGAAGACAGAGCCCGGCAAGAGGAAGAGCGCAGAAGGCAGGAGGAGGAGCGCACAAAGCGCGATGCAGAAGAGAAGAGGCGGCAAGAGGAAGGCTACTATTCTCGCTTGGAGGCTGAACGGAGGCGACAGCACGATGAGGCCGCCCGCCGCCTGTTGGAACCAGAGGCACCGGGCCTTTGCCGCCCACCGCTTCCCCGAGACTATGAGCCCCCCAGTCCTAGCCCTGCACCCGGCGCACCTCCTCCCCCCCCACAAAGGAATGCCAGCTACTTGAAGACGCAAGTGTTGAGCCCTGATTCCCTTTTCACCGCTAAATTCGTGGCCTATAATGAGGAGGAGGAGGAGGAGGATTGTAGTCTCGCTGGCCCCAACTCTTATCCGGGCTCTACCGGTGCAGCAGTCGGAGCCCACGATGCATGTAGGGACGCCAAAGAGAAGCGCAGCAAGTCCCAAGACGCCGACTCCCCCGGCAGTTCTGGTGCCCCCGAAAACCTGACCTTTAAGGAGCGCCAACGCCTGTTCTCACAGGGGCAGGACGTGAGTAACAAGGTAAAGGCATCCAGAAAGCTGACCGAGCTGGAAAACGAGCTCAACACCAAA
